# BoolSim, a Graphical Interface for Open Access Boolean Network Simulations and Use in Guard Cell CO_2_ Signaling

**DOI:** 10.1101/2021.03.05.434139

**Authors:** Aravind Karanam, David He, Po-Kai Hsu, Sebastian Schulze, Guillaume Dubeaux, Richa Karmakar, Julian I. Schroeder, Wouter-Jan Rappel

**Affiliations:** Physics Department, University of California, San Diego, La Jolla, CA 92093, USA; Division of Biological Sciences, Section of Cell and Developmental Biology, University of California, San Diego, La Jolla, CA 92093-0116, USA

## Abstract

Signaling networks are at the heart of almost all biological processes. Most of these networks contain a large number of components and often the connections between these components are either not known, or the rate equations that govern the dynamics of soluble signaling components are not quantified. This uncertainty in network topology and parameters can make it challenging to formulate detailed mathematical models. Boolean networks, in which all components are either on or off, have emerged as viable alternatives to more detailed mathematical models but can be difficult to implement. Therefore, open source format of such models for community use is desirable. Here we present BoolSim, a freely available graphical user interface (GUI) that allows users to easily construct and analyze Boolean networks. BoolSim can be applied to any Boolean network. We demonstrate BoolSim’s application using a previously published network for abscisic acid-driven stomatal closure in *Arabidopsis.* We also show how BoolSim can be used to generate testable predictions by extending the network to include CO_2_ regulation of stomatal movements. Predictions of the model were experimentally tested and the model was iteratively modified based on experiments showing that ABA closes stomata even at near zero CO_2_ concentrations (1.5 ppm CO_2_).

**One Sentence Summary:** This study presents an open-source, graphical interface for the simulation of Boolean networks and applies it to an abscisic acid signaling network in guard cells, extended to include input from CO_2_.

## Introduction

Intra-cellular signaling networks are essential in almost all biological processes. These networks are often complex, involving a large number of components (or nodes) that are inter-connected. To gain insights into these networks, it is possible to construct mathematical models. One of the strengths of these mathematical models is the ability to develop predictive outcomes of experimental perturbations (Phillips, 2015, Shou et al., 2015). These perturbations can be much more easily implemented in simulations than in experiments. Removing or changing a component or connection between components is a trivial task in simulations but usually is a task that requires lengthy wet lab experimental procedures. Predictions developed through models can enable narrowing the parameters for subsequent wet lab examination. Wet lab examination, in turn, can be used to iteratively update and correct mathematical models. Furthermore, mathematical models can be used to test potential biological mechanisms or can be utilized to pinpoint the most important components of a signaling network (Brodland).

One way of constructing mathematical models for signaling networks is to create a rate-equation model. In such a model, the concentrations for network components can take on all real values, and their change is governed by differential equations involving rate constants and the concentration of diverse components (Melke et al., 2006, Muraro et al., 2011, Wang et al., 2017, Hills et al., 2012). Such models, while useful, can quickly cease to be analytically tractable as the number of components increases. Furthermore, the concentrations of soluble signaling components are not well-defined and the strength of many of the connections between different nodes of the network is either poorly understood or not known at all. This results in considerable uncertainty in parameter values, especially the rate constants, potentially limiting the value of these mathematical models. This is particularly the case for signal transduction networks of soluble components for which rate constants are more difficult to define in a cellular context than, for example, metabolic flux networks (Feist et al., 2007).

An alternative to these “analog” networks is to formulate the problem in terms of Boolean networks (Kauffman, 1969). In these binary networks, each node can be either “ON” (1, or high) or “OFF” (0, or low) (Bornholdt, 2008, Wang et al., 2012, Schwab et al., 2020). The state of the nodes is then determined by an update rule, which involves information from the upstream nodes. In Boolean networks, the regulation is no longer encoded in terms of rate constants, which may or may not be quantified by experiments, but in terms of NOT, AND, and OR logic gates. For example, in the case of an AND gate, a downstream node will be turned on (i.e., 0 transitions to 1) if and only if the upstream node is on.

Despite the significant simplification associated with the binarization, Boolean networks have been shown to be able to predict behavior in a wide variety of networks, including genetic networks (Kauffman, 1969, Herrmann et al., 2012), protein networks (Dahlhaus et al., 2016), synthetic gene networks (Zhang et al., 2014), and cellular regulatory networks (Li et al., 2004, Lau et al., 2007, Albert et al., 2017). The price one has to pay for the simplification, however, is the loss of dynamic information. Moreover, the order in which the update rules are applied can critically affect the outcome of the network. Therefore, the networks are mainly used to probe steady-state conditions for the network.

In plants, Boolean networks have been applied to genetic networks to investigate, for example, possible crosstalk and microbe response (Genoud and Métraux, 1999, Timmermann et al., 2020). Boolean logic has also extensively been used to study abscisic acid (ABA)-induced stomatal closure in *Arabidopsis thaliana* (Li et al., 2006, Waidyarathne and Samarasinghe, 2018, Maheshwari et al., 2020, Maheshwari et al., 2019, Albert et al., 2017). This ABA signal transduction network contains a large number of components (>80), with many unknown rate constants, and is thus challenging to encode using an analog model. In a series of papers, it was shown that the formulation of a Boolean network for ABA-induced stomatal closure was able to confirm interesting experimental data (Albert et al., 2017, Maheshwari et al., 2020, Maheshwari et al., 2019). Furthermore, it was shown that the Boolean network could function as a vehicle to generate predictions. Specifically, predictions were generated through perturbations that either removed nodes or set nodes permanently to the “on” state. Some of these predictions were subsequently tested and validated in novel quantitative experiments (Albert et al., 2017, Maheshwari et al., 2019).

The aforementioned studies have clearly demonstrated the potential value of casting signaling pathways into Boolean networks. However, encoding these networks, especially ones with a large number of components, might present a significant impediment to wider spread use of Boolean networks to probe, analyze, and understand signaling networks. Motivated by the challenge of creating Boolean networks that can be used by the community for independent simulations, we present in this paper an open-source, user-friendly algorithm that can simulate Boolean networks that can be easily formulated by the users. This algorithm, which we term

BoolSim, uses a graphical user interface (GUI) and allows the users to define nodes and their internode connections, add nodes, subtract nodes, introduce mutations, and analyze the results. BoolSim can be freely downloaded from the GitHub repository (https://github.com/dyhe-2000/BoolSim-GUI or https://github.com/Rappel-lab/BoolSim-GUI). User friendly instructions for download and using BoolSim are provided in the Supplementary Text. Although BoolSim can be applied to any Boolean network, we first verify its use using the abscisic acid signaling network described by Albert et al. (Albert et al., 2017). We then describe an extension of this network that incorporates input from CO_2_ and examine ABA-induced stomatal closing, while clamping CO_2_ concentration in the leaf to very low levels (Zhang et al., 2018, Raschke, 1975). Finally, we test predictions from this extended network using quantitative experiments, and results from these experiments were used to update possible CO_2_ input mechanisms into the network.

## RESULTS

### Algorithm

Our algorithm, BoolSim, simulates a Boolean network with user-defined variables and interactions. Each node of the network corresponds to a Boolean variable that can be in one of two states: 0 (OFF) or 1 (ON). In biophysical terms, an ON state of a node corresponds to a concentration of its active form that is high enough to affect change through its interactions in the system. Conversely, an OFF state corresponds to a low concentration, not able to affect change. Nodes can flip their states as a result of their interactions with other nodes and these interactions are encoded using an update equation. Some nodes, denoting external conditions or inputs, remain fixed but affect the states of other nodes.

An update equation for a given node relates its future state to the current states of all upstream nodes. While there are several ways of formulating an update equation, our algorithm uses the so-called Sum of Products form that is intuitive and easy to formulate (Kime and Mano, 2003). This form consists of a series of terms linked by the OR logic operator. Each of these terms is a product, which contains Boolean variables and/or their negations connected by the AND operator. A typical example, used here to exemplify Boolean logic and extracted from the model of Albert and colleagues (Albert et al., 2017), reads:

AnionEM = SLAC1 | SLAH3 & QUAC1

where the symbols | and & stand for OR and AND, respectively. In this example, the future state of node AnionEM, which represents the anion efflux through the plasma membrane, is determined by the current states of three other nodes that encode two major classes of anion channels in guard cells: SLAC1, SLAH3, and QUAC1. Specifically, AnionEM is set to 1 or remains 1 if either a) SLAC1 is 1, or b) both SLAH3 and QUAC1 are 1 (Albert et al., 2017). If neither condition is met, AnionEM remains 0 or is set to 0. The Supplementary Text has a brief primer on Boolean algebra and several illustrations for formulating Boolean equations using the Sum of Products form.

In a typical simulation in BoolSim, there are a few nodes, which affect the nodes downstream of them but do not have any nodes upstream. As such, their states remain unchanged during the simulation. These nodes are called “input” nodes. There is typically one “output” node denoting the product or end state of the pathway, which can be monitored to determine the function of the network. For example, in the case of the ABA signaling network, ABA, Nitrite, GTP, etc., are inputs and the output node, termed “Closure”, represents stomatal closure (Albert et al., 2017). The initial states of other network nodes can be specified, based on prior knowledge, or can be assigned a random (0 or 1) value.

Once the initial states of all the nodes are specified, the program evaluates the update equations in a random order and exactly once for each node, with the output node evaluated last. The update for a single node is based on the current state of the network so that some of its upstream nodes may already have been updated in the same time step. This so-called asynchronous updating is motivated by the fact that many of the reaction rates are unknown, resulting in non-deterministic outcomes. A single time step in the simulation corresponds to each node being updated once. The number of time steps, corresponding to one iteration over all the nodes, can be chosen by the user and should be large enough for the system to reach a steady state. Furthermore, the user also specifies the number of simulations, each with randomly chosen initial conditions. All parameters, equations, and initial conditions can be easily entered into BoolSim using an intuitive graphical interface. Finally, BoolSim is able to display graphs of the dynamics of one or more nodes and all variables are stored for later analysis.

### Simulations Simple example

In a first example, we investigate a very simple Boolean network, shown in Fig. 1A. This network contains an input node (IN), an output node (OUT), and three intermediate nodes X, Y, and Z. The input node does not depend on any other nodes, which is simply written as IN = IN. Furthermore, X and Y reinforce each other and X inhibits Z through a NOT link (denoted by the symbol ∼) while IN inhibits X, Y inhibits OUT and Z activates OUT. For both X and OUT, the two inputs combine according to AND logic as denoted. The Boolean update equations can thus be simply written as follows:

**Figure 1:**
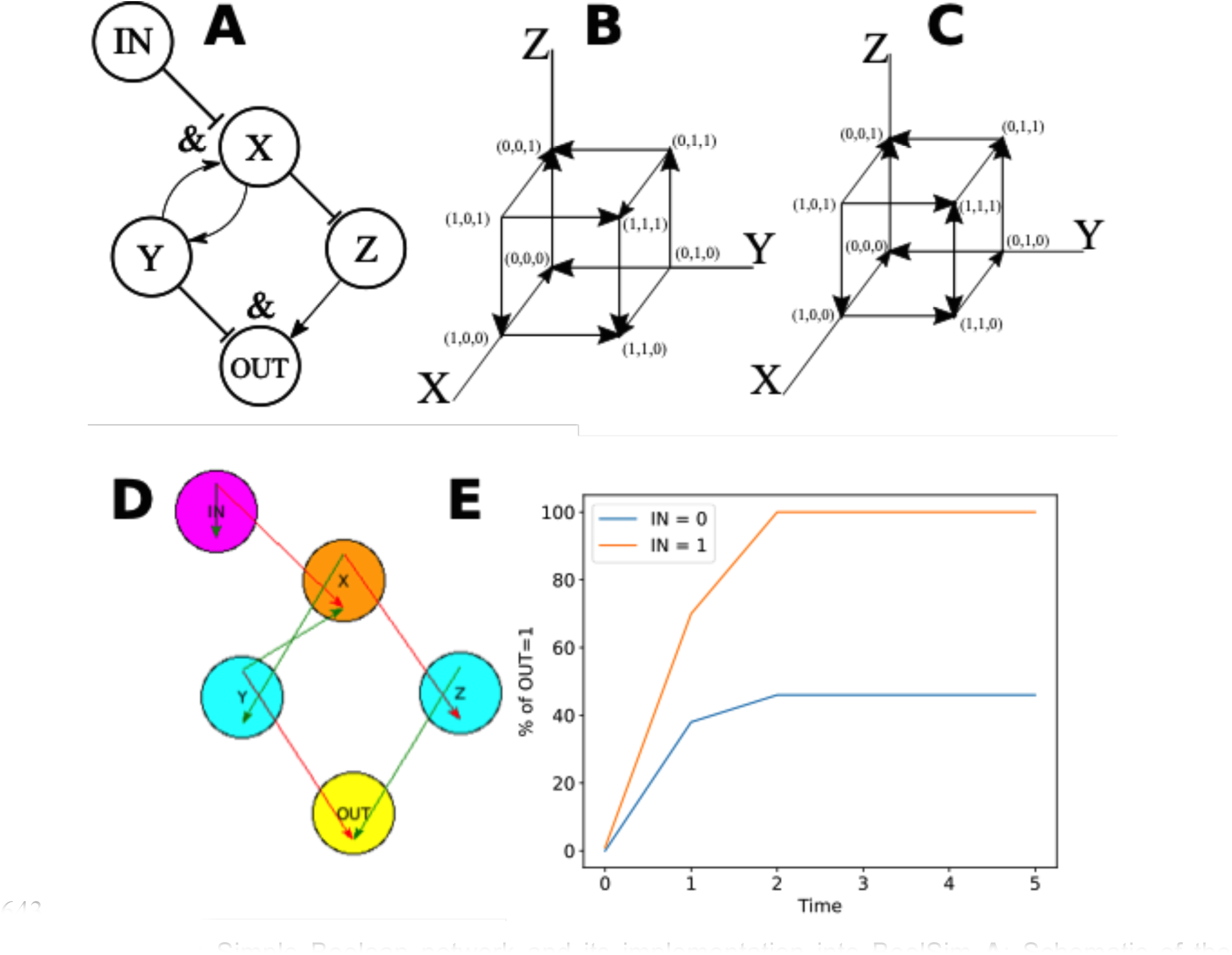
Simple Boolean network and its implementation into BoolSim **A**: Schematic of the network. Pointed arrowheads indicate positive regulation and flat arrowheads indicate negative regulation. For nodes X and OUT, the two inputs combine using AND logic, indicated by the symbol &. **B:** State space and dynamics, represented by arrows, in the absence of input (IN = 0). The two possible attractors, (0,0,1) and (1,1,0), are indicated by red dots. **C:** State space and dynamics in the presence of input (IN = 1). The only one attractor, (0,0,1), is indicated by a red dot and leads to OUT = 1. **D**: Representation of the Boolean model in the BoolSim GUI. Green and red arrows indicate positive and negative regulation, respectively. An arrow within the node indicates self-regulation, which can be positive or negative. In this representation, one can ‘search’ for upstream and downstream nodes of a given node. Here, X (colored in orange) has two downstream nodes, colored in cyan and one upstream node, colored in magenta. Other node(s) are colored in yellow. **E**: Percentage of OUT=1 vs. time in an asynchronous update scheme starting with randomly assigned initial states for the intermediate nodes when the input is present (IN = 1, orange line) and absent (IN = 0, blue line).

IN = IN

X = Y & ∼IN Y = X

Z = ∼X

OUT = ∼Y & Z

The dynamics of this network can be worked out by hand or, since it only contains 3 nodes, can be analyzed graphically. One feature of an asynchronous update scheme is that the updated state is always the ‘nearest neighbor’ of the previous state. Therefore, the evolution of the states can be represented as a continuous trajectory through the state space with dimensions equal to the number of nodes. This is shown in Fig. 1B&C for our simple model in the absence (IN = 0; B) and in the presence of input (IN = 1; C). The 3-dimensional state space is spanned by X, Y, and Z and all possible states of the intermediate nodes are points in this 3-dimensional space that are connected by arrows according to the rules of the Boolean network. By following these arrows, the attractors for the model can be determined. For example, in the absence of input (Fig. 1B), the state X=1, Y=0, and Z=0, compactly written as (1,0,0) (Fig. 1B), can either transition to (0,0,0) or to (1,1,0). Since no arrows originate from (1,1,0), this is an attractor of the system: this state will remain unchanged indefinitely. Furthermore, the only permissible transition from (0,0,0) is to (0,0,1), which can easily be seen to be an attractor as well. Since only (0,0,1) leads to the ON state of the OUT node, we find that OUT=1 in 50% of the possible initial conditions. In the presence of input (IN=1), however, it is easy to verify that the only possible attractor is (0,0,1) and thus OUT = 1 for all initial conditions. Calculations and further analysis on this simple network are provided in the Supplementary Text.

The implementation in BoolSim is shown in Fig. 1D, obtained after specifying the input files in the subfolder “sample_data_files/simple_network_data_files” of the repository (https://github.com/dyhe-2000/BoolSim-GUI or https://github.com/Rappel-lab/BoolSim-GUI). Here, the green/red arrows indicate positive/negative regulation and an arrow within the node indicates self-regulation (green/red for positive/negative). In BoolSim, all nodes are visualized in yellow by default. Connections between a particular node and other network nodes, however, can be easily visualized by double-clicking on the specific node, after which it changes its color to orange. Upstream nodes will then change their color to magenta while downstream nodes will turn cyan. In the example of Fig. 1D, this procedure has been carried out for node X. Note that the color scheme for the nodes can be changed by the user (see Supplementary Text section 2, Downloading and Running BoolSim).

This network was simulated using BoolSim, choosing 5 time steps and 50 different sets of initial conditions. The results of these simulations are shown in figure 1E, where we plot the state of the output node, expressed as the percentage of runs in which OUT=1, in the absence (blue line) and presence (orange line) of input as a function of time. Note that time in these plots indicates the iteration number. The state of the simulation that is of physiological relevance is the steady state, here reached after two steps. Consistent with the arguments presented above, the simulations show that this percentage is 50% when IN=0 and 100% when IN=1.

### ABA network

Next, we applied BoolSim to the ABA-induced stomatal closure network formulated by Albert *et al*. (Albert et al., 2017). The input file for this network, containing all components by name and their interactions, can be found in the subfolder “sample_data_files/ ABA_data_files/” of the repository for the Boolean equations and for the names of the nodes (https://github.com/dyhe-2000/BoolSim-GUI or https://github.com/Rappel-lab/BoolSim-GUI). The reconstruction of the published ABA signaling network (Albert et al., 2017) within the BoolSim interface here, will enable any user to use and manipulate components of this network and develop experimental predictions, as well as modify the Boolean network depending on experimental outcomes or to predict outcomes for modified network models. A screenshot of our implemented network encoded within the BoolSim GUI is presented in Fig. 2, with the input ABA node shown in red and the “Closure” output node shown in green. This network contains 81 nodes, including input and output nodes, and was constructed by and adapted from Albert *et al*. (Albert et al., 2017) following an extensive survey of more than one hundred peer-reviewed articles. As in the simple example, the interactions between nodes in the GUI are color-coded, with green arrows representing positive interactions and red arrows representing negative interactions. Using the BoolSim GUI interface, the user can move nodes around by simply dragging them to a new location. Furthermore, to facilitate examining inter-node connections, double-clicking on a node reveals all downstream and upstream interactions of that node. (see https://github.com/dyhe-2000/BoolSim-GUI or https://github.com/Rappel-lab/BoolSim-GUI; Detailed instructions can be found in the Supplementary Text.)

**Figure 2:**
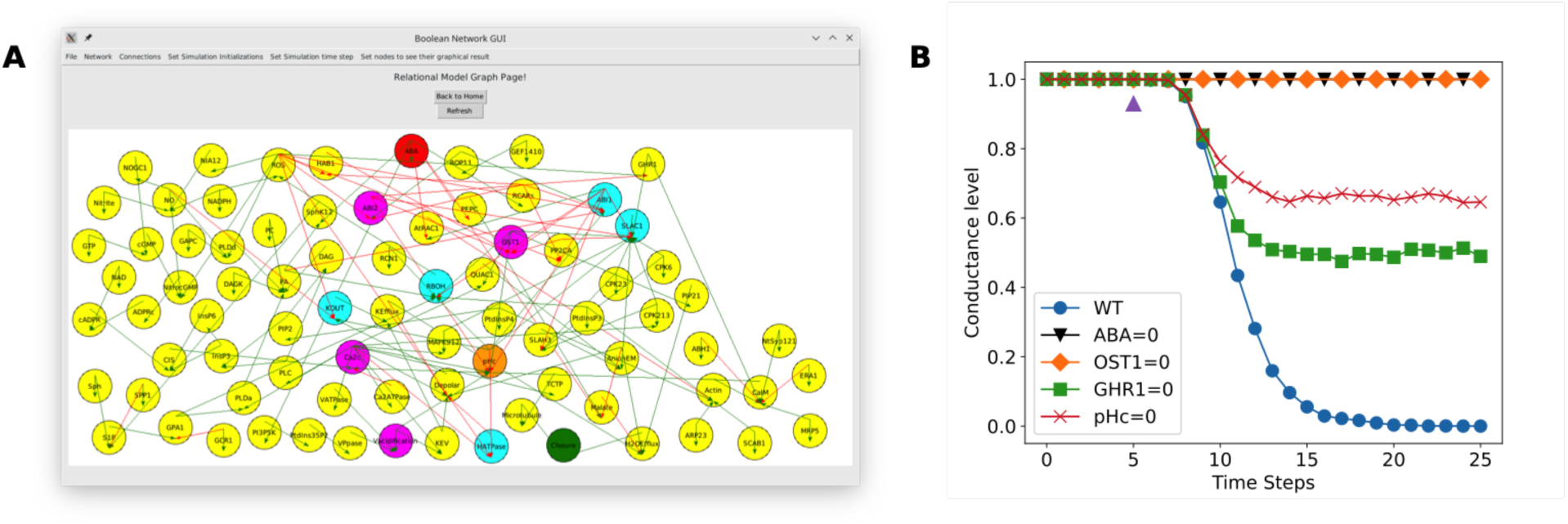
Implementation of ABA into BoolSim **A:** Visualization of the Boolean network for ABA- induced stomatal closure, rendered here by the BoolSim GUI. The input ABA and output Stomatal Closure are colored in red and green, respectively. The node denoting cytoplasmic pH (pHc), colored in orange, has its upstream nodes colored in magenta and downstream nodes in cyan. Connections for any node can be viewed by double-clicking on the node of interest (see Results). **B:** Stomatal conductance as a function of time steps in the simulation for wild type and the mutants *ost1* and *ghr1*, and alteration of cytosolic pH (pHc) (see Results). A conductance level of 1 corresponds to maximal relative stomatal conductance while 0 represents complete stomatal closure. The violet triangle shows the point in the simulation where ABA is introduced (except for the case ABA=0).

Results of the BoolSim simulations for 25 time steps and averaged over 2,500 initial conditions are presented in Fig. 2B. In this, and subsequent curves, we chose to illustrate the predicted stomatal conductance level, computed as 1-Closure, as a function of time step. Reporting the conductance level, which varies between 0 (corresponding to closed stomata) and 1 (corresponding to open stomata), facilitates comparison with experiments in which the stomatal conductance is presented (e.g., Fig. 4). In the absence of ABA, simulated by setting the input node ABA to 0 throughout the simulation, the output node Closure is 0 for all time steps, corresponding to no stomatal closure and a conductance level of 1 (orange diamond curve). In the presence of ABA, modeled by changing ABA from 0 to 1 at time step 5, the network reaches a conductance level of 0 (stomatal closure) after approximately 15 time steps (blue curve, labeled as wild type (WT)). By implementing the model of Albert *et al*. (Albert et al., 2017), we have also computed the relative stomatal conductance for several mutants, labeled in Fig. 2B, following the introduction of ABA after 5 time steps. Knocking out the Open Stomata 1 protein kinase (OST1), which corresponds to forcing the node OST1 to 0 at all times, results in a conductance level that remains 1 (no stomatal closure) after setting ABA=1 (orange curve) Furthermore, knocking out GHR1 (Guard cell hydrogen peroxide resistant 1) results in a conductance level of 0.5 (50% stomatal closure) following the introduction of ABA (green curve) and manipulating the cytosolic pH through the node pHc leads to a conductance level of approximately 0.65 (red curve). These control predictions are identical to the ones obtained in the original publication (Albert et al., 2017), further validating reconstruction of this network in our interactive graphical user interface.

### CO_2_ network

To illustrate how one could use our GUI interface to examine and explore networks, we have taken the original ABA network of *Albert et al.* and have first extended it with a putative branch that models the input of carbon dioxide (CO_2_). Elevated CO_2_ triggers stomatal closure and some elements of CO_2_ signaling overlap with those of ABA signaling, whereas others affect stomatal closure through separate pathways (Merilo et al., 2015, Hsu et al., 2018, Zhang et al., 2020). Based on previous experimental data, we modeled the CO_2_ branch to be upstream of GHR1 in the ABA network (Hõrak et al., 2016, Jakobson et al., 2016). The added branch contains CO_2_ as input, which is then catalyzed by the beta carbonic anhydrases βCA4 and βCA1 in parallel (Hu et al., 2010, Hu et al., 2015). These then activate the node MPK12/MPK4 via yet unknown mechanisms. This node inhibits the negative-regulator of CO_2_-induced stomatal closing HT1, which, in turn, is regulates CBC1/CBC2 either directly or indirectly (Hashimoto et al., 2006, Hõrak et al., 2016, Hiyama et al., 2017). Finally, CBC1/CBC2 enters the ABA network through an assumed inhibitory link to GHR1 (Fig. 3A).

**Figure 3:**
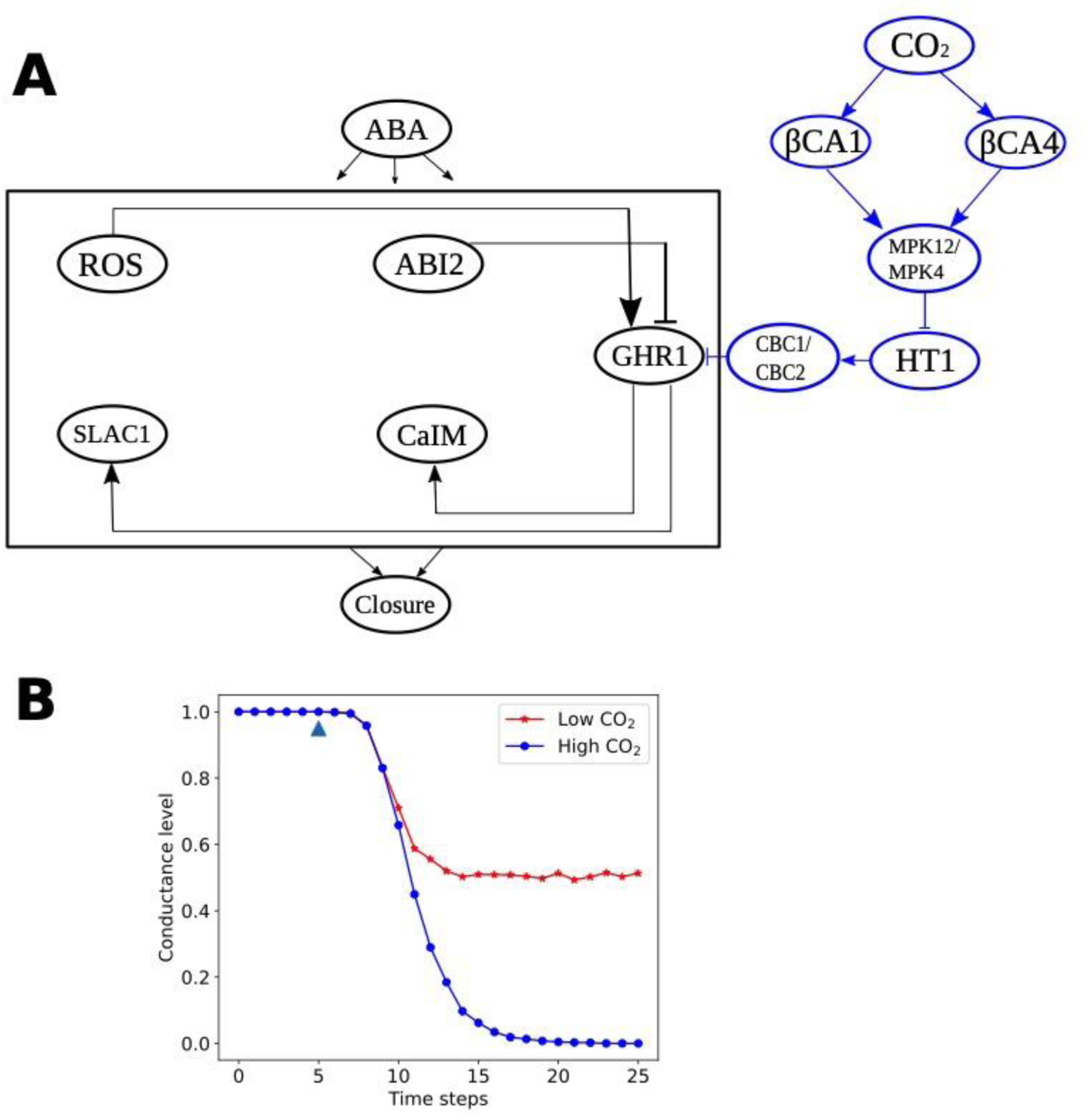
Stomata 2.0 and BoolSim **A**: ABA-driven stomatal closure model extended with a CO_2_ branch, indicated in blue, which positively regulates GHR1. The box denotes all the intermediate nodes of the original ABA network shown in Fig. 2 with only GHR1 and its immediate upstream and downstream nodes shown. **B**: Predicted relative stomatal conductance levels obtained by implementing Stomata 2.0 into BoolSim for two concentration levels of CO_2_: low (CO_2_=0; red line and symbols) and high (CO_2_=1; blue line and symbols). The blue triangle shows the point in the simulation where ABA is introduced (ABA=1). Without the introduction of ABA (ABA=0) and independent of CO_2_, conductance remains high and stomata do not close.

The above summarized CO_2_ branch can be translated into the following Boolean equations:

CO_2_ = CO_2_

βCA1= CO_2_

βCA4= CO_2_

MPK12/MPK14 = βCA4| βCA1

HT1 = ∼ MPK12/MPK14

CBC1/CBC2 = HT1

GHR1 = ∼ ABI2 & ROS & ∼ CBC1/CBC2

Note that the equation for GHR1 is adapted from Albert et al and takes into account the existing connections from the ABA network (from ABI2 and ROS) and the new input from the CO_2_ branch. This simplified CO_2_ signaling model, which can be accessed in the online BoolSim repository, termed as Stomata 2.0, includes presently identified and confirmed early CO_2_ signaling mechanisms that have been found to function in the CO_2_ signaling pathway upstream of the merging with the ABA-induced stomatal closing pathway (Hsu et al., 2018, Zhang et al., 2018, Zhang et al., 2020).

Based on this model, we then determined how the extended network responds in simulations to ABA under high and low CO_2_ conditions (Fig. 3B), simulated by setting the input node for CO_2_ to either 0 (very low concentration) or 1 (high concentration). For CO_2_=1, the introduction of ABA at time step 5 results in a decrease of conductance level from 1 to 0 (Fig. 3B, blue curve), identical to the WT response shown in Fig. 2 (blue curve). This can be understood by realizing that the added CO_2_ branch has no effect on the ABA network since CBC1/CBC2 is 0 when CO_2_=1. When simulating very low CO_2_ conditions, our simulations predicted that introduction of ABA also induces stomatal closure and, thus, a decrease in the conductance level. The conductance level in the presence of low CO_2_, however, was found to be only reduced from 1 to 0.5 (Fig. 3B, red curve).

To test our predictions experimentally, we analyzed ABA-mediated stomatal closure under either 400 ppm or near 0 ppm (∼1.5 ppm) [CO_2_] by conducting gas-exchange experiments with ABA application to the transpiration stream of excised intact leaves (Ceciliato et al., 2019). Our results show that application of 2 µM ABA induced robust stomatal closure in leaves exposed to 400 ppm [CO_2_] as expected (blue curves Fig. 4A-D). As stomatal responses are known to show biological noise, and as ABA-induced stomatal closing has not been previously analyzed at near zero CO_2_, we conducted four independent sets of experiments (Fig 4). In all four experiments, leaves exposed to 1.5 ppm [CO_2_] showed stomatal closing in response to 2 µM ABA, with a degree of expected biological variation (red curves Fig. 4A-D). By analyzing the steady state stomatal conductance in leaves it appears that the response to ABA at low CO_2_ was reduced in 3 of these experiments (Fig. 4A, C&D). We also compared the steady-state stomatal conductance before and after applying ABA (Fig. 4E-H). This analysis shows that independent of whether leaves are exposed to 400 ppm CO2 or 1.5-2 ppm CO2, the ABA responses had a similar magnitude in three of the experiments and a stronger ABA response in one of the experiments (Fig. 4F). Furthermore, our data show that in the absence of ABA, leaves exposed to low CO_2_ show a higher stomatal conductance than leaves exposed to 400 ppm CO_2_. This is consistent with a reduction in stomatal conductance upon CO_2_ elevation. In addition, analyses of an early time point of the ABA response, 10 minutes after ABA addition, show a slightly, but significantly slowed initial ABA response in 3 of 4 experiments (Fig. E,G,H). Taken together, our experimental data show that the steady-state stomatal conductance responses to ABA remain to a large degree intact even at very low CO2 concentration, and indicate an approximately steady-state additive effect of low CO_2_ on the stomatal conductance prior to ABA exposure.

**Figure 4.**
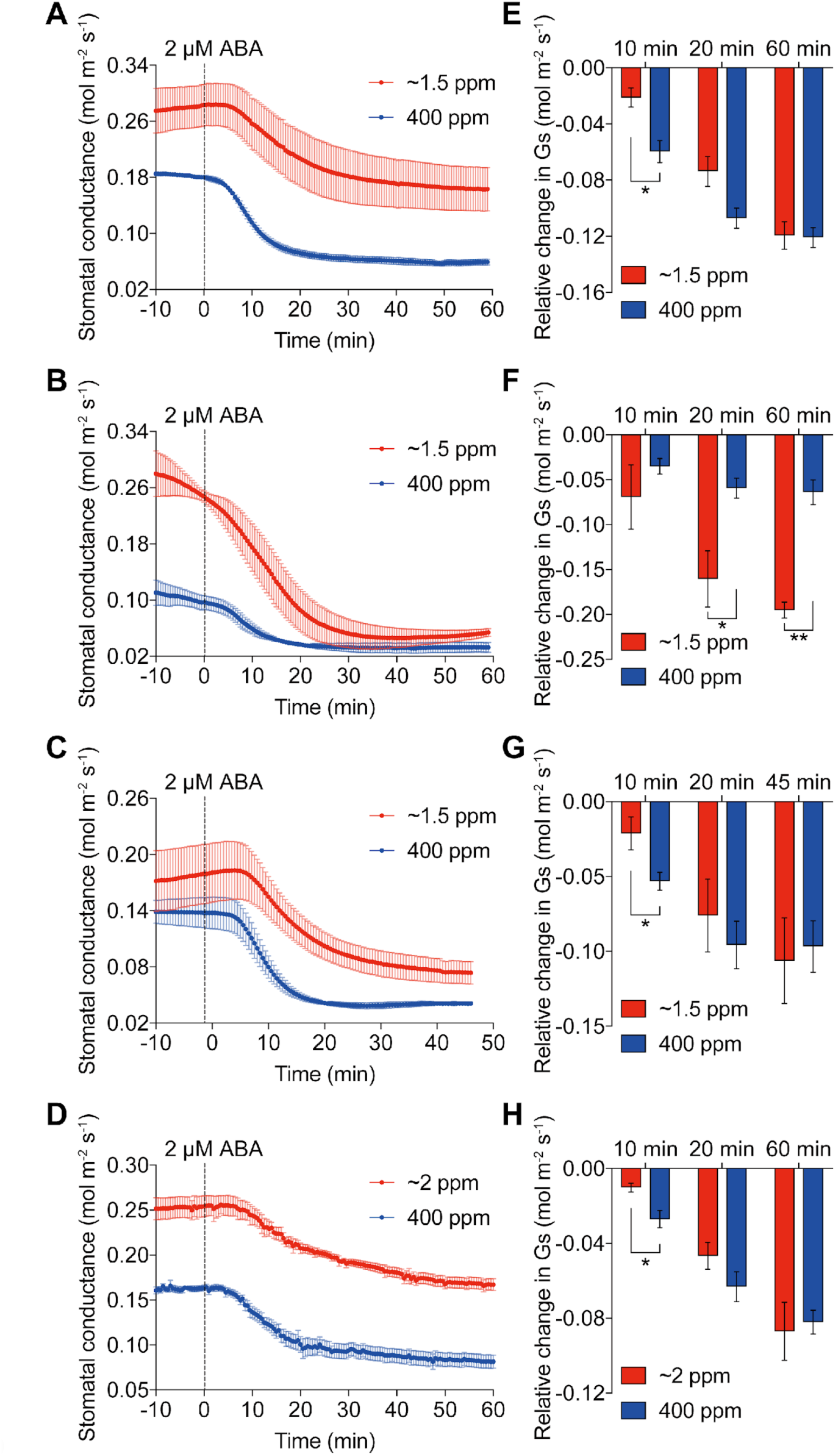
ABA-mediated stomatal closing responses of WT leaves during CO_2_ starvation. A to C, Intact excised leaves of wild-type plants (n = 3 to 4 independent leaves per treatment and experimental set) were equilibrated at 400 ppm CO_2_ or ∼1.5 ppm CO_2_ for 60 min prior to stomatal conductance measurements. Stomatal conductances were measured with the LI- 6400XT Portable Photosynthesis System. D, Time-resolved stomatal conductance responses to ABA in the intact excised leaves (n = 5 independent leaves per treatment) equilibrated at ∼2 ppm CO_2_ or 400 ppm CO_2_ for 2 hr prior to ABA application. Experiments were carried out using the LI-6800 Portable Photosynthesis System. In each experimental set, 2 µM ABA was applied through the transpiration stream via the petiole at time 0 min. CO_2_ concentrations in the intercellular spaces of leaves (Ci) equilibrated under CO_2_ starvation were computed using the gas exchange analyzer (see Methods), with Ci values < 20 ppm (A to C) and < 3 ppm (D) before application of ABA. E to H, Relative changes in stomatal conductance at the indicated time points after ABA application compared to 0 min. Data present mean ± SEM. * P < 0.05 and ** P < 0.01 student’s t-test in E to H.

A comparison of our experimental and model results in the absence of ABA reveals that Stomata 2.0 (Fig. 3B) is not able to fully capture the dependence of the steady-state conductance levels on CO_2_. This suggests that additional modifications of the ABA network are required to reproduce the observed reduction of stomatal conductance in the presence of CO_2_.

To explore possible modifications that result in model predictions that are more consistent with our experimental steady-state response results, we utilized the ability of BoolSim to easily modify, simulate, and visualize modified Boolean network outcomes. We found that we were able to better reproduce experimental data if we modified the Boolean equations for four network components (Fig. 5A). Specifically, we modified the nodes Ca2c (cytosolic calcium), which is linked in the ABA model (Albert et al., 2017) to CaIM (Ca2+ influx across the plasma membrane), Microtubule (Microtubule depolymerization) and H2OEfflux (water efflux through the plasma membrane) to:

**Figure 5:**
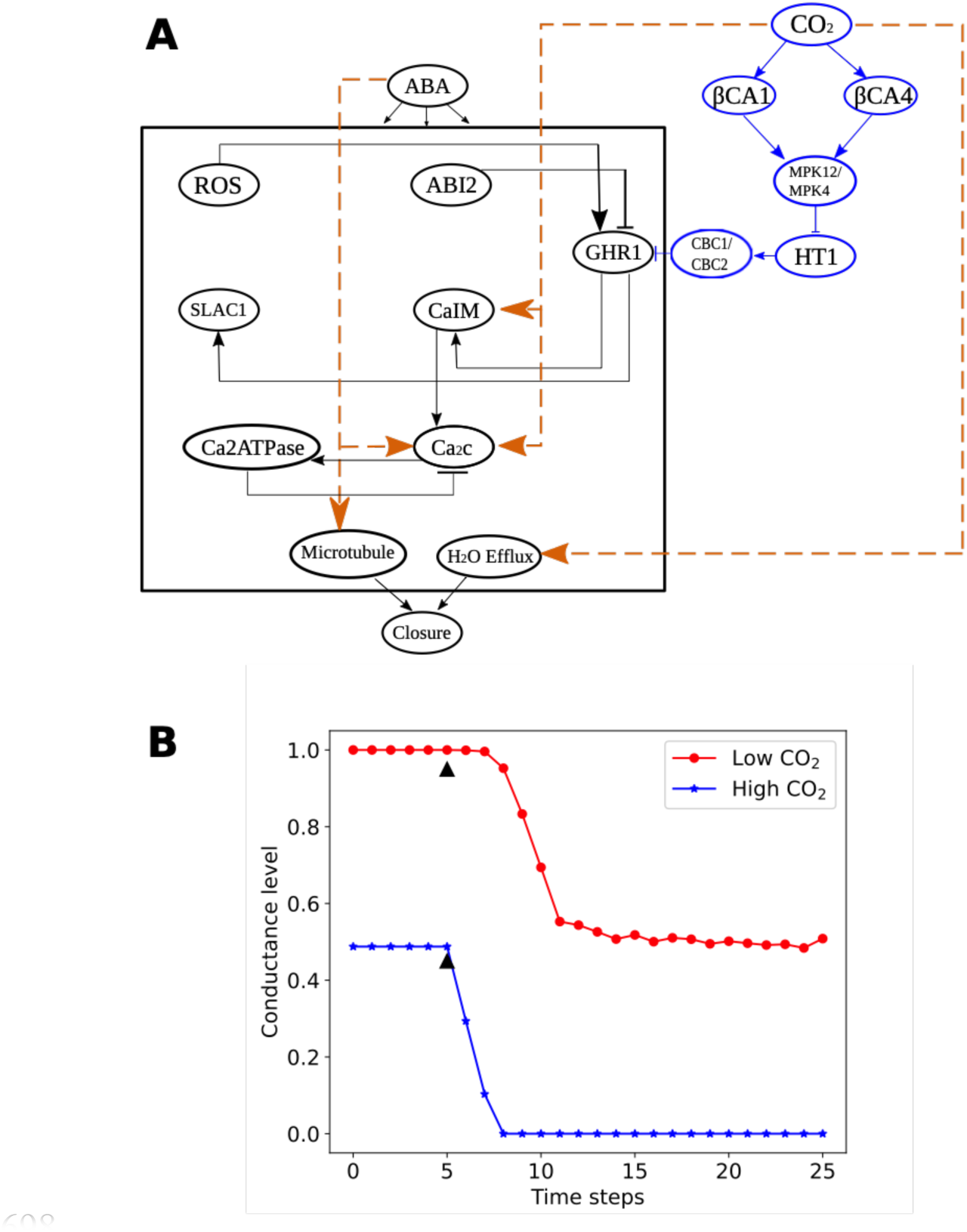
Stomata 2.1 and BoolSim **A**: ABA-driven stomatal closure model extended with a CO_2_ branch, indicated in blue, which positively regulates GHR1, and additional modifications represented by the orange links (see Text for details). **B-C**: Predicted relative stomatal conductance levels obtained by implementing Stomata 2.1 into BoolSim for two concentration levels of CO_2_: low (CO_2_=0; red line and symbols) and high (CO_2_=1; blue line and symbols). The red triangle shows the point in the simulation where ABA is introduced.

Ca2c = ∼Ca2ATPase & (CIS | CaIM) <u>| (ABA&CO2)</u>

CaIM = ∼ABH1 & (NtSyp121 | MRP5 | GHR1) | ∼ERA1 | Actin <u>| CO2</u>

Microtubule = TCTP | Microtubule <u>& ABA</u>

H2OEfflux = AnionEM & PIP21 & KEfflux & ∼Malate <u>| CO2</u>

where the modifications are underlined. Introducing a CO_2_ dependence on calcium signaling was motivated by experimental evidence that cytosolic calcium involved in CO_2_-induced stomatal closure (Schwartz et al., 1988, Webb et al., 1996, Schulze et al., 2021). Furthermore, findings that anion efflux and water efflux functions in the CO2 response and that GHR1 functions in CO_2_-induced stomatal closure were included (Hõrak et al., 2016, Jakobson et al., 2016). Addition of an ABA component to microtubule function was motivated based on recent findings (Qu et al., 2018, Rui and Anderson, 2016). These modifications, their associated Boolean logic operations, and their effect on closure are further detailed in the Supplementary Text. When simulating this modified and updated network, termed Stomata 2.1, we find that the steady-state conductance level now depends on CO_2_ and is reduced for high CO_2_ conditions (Fig. 5B). Furthermore, and also consistent with the experiments, the introduction of ABA reduces stomatal conductance by similar absolute conductance changes for both low and high CO_2_ conditions (Fig. 5B). The Stomata 2.1 network is able to better reproduce experimental results. We have included the network files for both Stomata 2.0 and 2.1 in the folder sample_data_files/ in the BoolSim repositories. We anticipate that community members will be able to use BoolSim as a starting point to easily introduce modifications and iteratively test predictions.

## Discussion

Large signaling networks are common in biology in general, and plant physiology. The published ABA signaling network we implemented into BoolSim, for example, contains more than 80 components (Albert et al., 2017). The vast majority of the interaction strengths between these components is not known, making it difficult to formulate mathematical models of these networks. Motivated by the simplicity and utility of Boolean networks and the challenges associated with formulating detailed rate equation-based models for these large networks, we have presented here a software package with a graphical user interface (GUI) that can simulate, visualize, and plot the results of a user-defined Boolean network. Our package, named BoolSim, is free to use and distribute, and is built from free and open-source software. The interface is intuitive and users do not require extensive coding knowledge to use it. The Supplementary Text contains detailed instructions on downloading, installing, and running BoolSim on Windows, Mac, and Linux-based machines.

BoolSim uses asynchronous and random order of update, which is best suited to simulate a network of chemical reactions in which the outcome of one reaction then affects the outcome of another in the near future (hence asynchronous), and when the relative rates of different reactions in the network are unknown (hence random order of update). Besides nodes and connections, users may also specify the number of time steps or iterations to run the simulations and the number of initial conditions to get a statistically robust sample. Once a system is defined, it may be visualized as a network in BoolSim. Visualization includes identifying the upstream and downstream nodes of a given node and the type of connections (activating or inhibiting) between them, by simply double-clicking on the node of interest. The steady states of nodes of the system after simulation can be quickly plotted within BoolSim. The trajectory of the entire simulation is stored in a NumPy array; a Jupyter notebook is provided with the package that can be used to further analyze the system starting from the NumPy array, including producing publication-ready plots of the simulation.

We first tested and verified BoolSim using a published advanced model for stomatal closure in guard cells as mediated by abscisic acid (ABA) and verified that our simulations were consistent with those of Albert *et al*. (Albert et al., 2017). We then extended the network to include the effects of CO_2_ on stomatal movements. Previous research indicated that CO_2_ might mediate signal transduction via the OST1 protein kinase, as *ost1* mutant leaves were impaired in their stomatal response to CO_2_ elevation (Xue et al., 2011, Merilo et al., 2013). However, more recent studies unexpectedly showed that, in contrast to abscisic acid, CO_2_ elevation does not activate the OST1 protein kinase(Hsu et al., 2018, Zhang et al., 2020). This research further provided experimental evidence that basal OST1 protein kinase activity and basal ABA signaling are required for WT-like CO_2_-induced stomatal closure (Hsu et al., 2018, Zhang et al., 2020) (Figs. 3 and 4). The biochemical link by which CO_2_ signaling merges with ABA signaling is thus proposed to lie downstream of the OST1 protein kinase, but remains unknown. In the present study, to test a simplified model merging the ABA signaling and CO2 signaling networks, we modeled this link to occur at the level of the transmembrane receptor-like (pseudo)kinase GHR1 (Hua et al., 2012, Sierla et al., 2018).

The simulations of this simplified network predicted that the response to ABA should depend on the CO_2_ concentration (Fig. 3). This prediction was then subsequently analyzed and ABA- mediated stomatal closure of intact leaves was measured while leaves were either exposed to ambient 400 ppm [CO_2_] or near zero (1.5 ppm) [CO_2_] (Fig. 4). Our data show that leaves exposed to 1.5 ppm [CO_2_] showed a robust response to ABA. Under low CO_2_ conditions the stomatal conductance remained higher prior to ABA application at steady state than at 400 ppm CO_2_ (Fig. 4A). When comparing true steady state stomatal conductance responses, it appears that CO_2_ and ABA may in part have additive responses (note that basal ABA signaling amplifies or accelerates the CO_2_ response (Hsu et al., 2018), such that the starting stomatal conductance was much higher at low CO_2_ due to the lack of CO_2_-induced stomatal conductance reduction (Fig. 4A).

In contrast to our experimental data (Fig. 4), Stomata 2.0 predicted identical conductance levels for low and high CO_2_ concentration in the absence of ABA (Fig. 3B). In an illustration of the use of BoolSim, we modified the ABA network further, with as goal to incorporate CO_2_ dependence on steady-state conductance levels when ABA is absent. Creating this updated network, Stomata 2.1, was greatly facilitated by the ability of BoolSim to easily implement changes and generate predictions. We introduced CO_2_ dependence on calcium signaling based on experimental evidence that cytosolic calcium is involved in CO_2_-induced stomatal closure (Schwartz et al., 1988, Webb et al., 1996, Schulze et al., 2021). Furthermore, it is well-established that anion efflux and water efflux from guard cells are essential for the CO_2_–induced reduction in stomatal conductance. Furthermore, findings that GHR1 functions in CO_2_-induced stomatal closure (Hõrak et al., 2016, Jakobson et al., 2016) were expanded to include GHR1 predictions of the original ABA signaling model (Albert et al., 2017). Addition of an ABA component to microtubule function was motivated based on recent findings on roles of guard cells microtubules (Qu et al., 2018, Rui and Anderson, 2016). The output of Stomata 2.1 was able to better incorporate effects of CO2 and the present experimental data (Fig. 5B). As described earlier, important gaps exist in the understanding of the CO_2_ signaling pathway, including that the primary CO2/bicarbonate sensors remain unknown in guard cells and the mechanisms by which HT1, CBC1 and CBC2 link to stomatal closing mechanisms is unknown. Expansion of the present model will be required.

Our proposed additions to the existing ABA network, which illustrate the potential use of BoolSim, are meant as a starting point for further explorations and further research is needed to determine the precise mechanism by which CO_2_ signaling merges with abscisic acid signal transduction. Nevertheless, several improvements can be suggested. First, it is conceivable that CO_2_ affects yet unknown mechanisms. Second, the CO_2_ pathway may contain feedback loops, which can be easily implemented within BoolSim. Finally, we should point out that Boolean networks do not incorporate explicit rate constants and contain nodes that can only take one of two values (0 or 1). Therefore, these networks are not able to address the kinetics of responses nor how they respond to graded inputs.

Most importantly, the GUI platform and stomatal signaling model developed here can be used and altered by users to test diverse predictions and to add expanded components or to or modify the ABA and CO_2_ signaling models. Thus, the methods and software tools presented here can be of interest to the wider plant biology community interested in physiological pathways and may be used to generate further predictions in the ABA- and CO_2_ signaling networks. Importantly, the BoolSim platform developed here can be applied to any Boolean network and can be used to generate publicly and modifiable Boolean models for any desired process.

## Materials and Methods

### Software

BoolSim is implemented using Python and C++ and can be freely downloaded from the GitHub repository (https://github.com/dyhe-2000/BoolSim-GUI and https://github.com/Rappel-lab/BoolSim-GUI). It requires a current version of Python and C++ and a detailed manual, including installation instructions, is provided in the GitHub repository. These instructions are provided for Windows, Mac, or Linux-based computers.

### Experiments

Plants of the *Arabidopsis thaliana* accession Columbia (Col-0) were grown as described in (Hsu et al., 2018). Stomatal conductance (Gs) measurements in response to ABA were performed in detached intact leaves of 5.5- to 7-week-old plants in *Arabidopsis* leaves following the procedure described previously (Ceciliato et al., 2019) using a LI-6800 Portable Photosynthesis System with an integrated Multiphase Flash Fluorometer (6800-01A; LI-COR Biosciences, Lincoln, NE, USA). Detached leaves were clamped in the leaf chamber and kept at ∼1.5 ppm or 400 ppm [CO_2_], 135 µmol m^-2^ s^-1^ red light combine with 15 µmol m^-2^ s^-1^ blue light, 70 ± 0.5% or 65 ± 0.5% relative air humidity, 21°C heat exchanger temperature, and 500 µmol·s^-1^ incoming air flow rate for at least 2 hour until stomatal conductance is equilibrated and stabilized. Stomatal conductances were recorded every 30 sec under ∼1.5 ppm or 400 ppm [CO_2_] for 10 min. ABA (2 µM) was then added to the transpiration stream via the petiole, and stomatal conductances were recorded as shown in the figure panels. In each independent set of experiments, 5 intact leaves from independent plants were analyzed per experimental condition.

## Acknowledgements

This research was funded by a grant from the National Science Foundation (MCB-1900567) to WJR and JIS. SS was supported by a Deutsche Forschungsgemeinschaft (DFG) fellowship (SCHU 3186/1-1:1). GD was supported by an EMBO long-term post-doctoral fellowship (ALTF334-2018).

## Supplementary Text

The following text contains instructions to download, install, and run BoolSim on a personal computer like an office laptop. BoolSim is free to run and distribute, and uses other open-source software packages including Python and C++. In addition, it describes the Boolean equations introduced in Stomata 2.1. BoolSim can be freely downloaded from the GitHub repository (https://github.com/dyhe-2000/BoolSim-GUI or https://github.com/Rappel-lab/BoolSim-GUI). This text contains detailed installation instructions for Windows-based machines while the repository contains instructions for the MacOS and Linux operating systems.

This document is structured as follows. Section 1 contains the installation instructions for C++ and Python for the Windows operating system. section 2 contains instructions to download BoolSim from the code-sharing website GitHub. Section 3 describes how to run a few example Boolean networks using BoolSim. Section 4 describes the process of designing your own Boolean network. It is advisable to try out the “simple network” (details in the main text and in Section 4) before running the ABA network or the updated Stomata 2.0. Creating a new Boolean network requires an elementary knowledge of Boolean algebra besides empirical data on the system one wishes to model. Section 5 contains a primer on Boolean algebra. In Section 6 we highlight some common errors encountered during installation and running of BoolSim and ways to get around them. Finally, Section 7 describes Stomata 2.1 and the modifications to the ABA network for CO_2_ signaling described in the main text.

### 1. Installing BoolSim - Software Requirements (Windows)

BoolSim requires that the GNU C++ compiler (called g++) and Python be installed on the user’s computer. Both the tools are freely available. The following steps run through the installation of the two programs.

1. Firstly, check if g++ is already installed on your computer. Open the command prompt by right-clicking on the start button and click on “command prompt”; alternatively, search for “command prompt” in the search bar in the start menu and open the application from the search results. Type ‘g++ --version’ without the quotes and hit Enter (see screenshot 1).

a. BoolSim requires the g++ version to be 6 or higher. If g++ is properly installed, the command prompt displays the version of g++ followed by a message. (See screenshot 2). Otherwise, it shows an error (see Section 6 for troubleshooting).
b. Once g++ is installed, we come back to this step to verify that the installation is proper and complete. If g++ is installed, please go to step #5.
2. To install g++ we first download and install a program called MinGW. This program installs the necessary compilers for us in a matter of a few clicks.
3. This video runs through g++ installation using MinGW (https://youtu.be/sXW2VLrQ3Bs). Follow the steps as shown in the video. Here are some tips you might find useful while working through the video:

a. URL to download MinGW: https://sourceforge.net/projects/mingw/
b. The installation of MinGW requires changing environment variables. The video explains one way of accomplishing this. A different way to change environment variables on your system is to open System Settings (which you can open by searching for it in the search bar in the start menu) and type ‘environment variables’ in the search bar of the system settings window.
4. After the g++ installation is complete, go to step 1. If successful, your computer can now compile and run c++ programs.
5. Next, we install Python. To check if Python is already installed, type ‘python --version’ in the command prompt and hit Enter.

a. As before, if Python is properly installed, the command prompt shows the version number and a message. If it is not installed, it displays an error (see Section 6 for troubleshooting).
b. BoolSim requires python3. Once Python is installed, we come back to this step to verify that the installation is proper and complete. If installed, go to step #7.
c. If python3 is installed on your computer, the above command shows a version 2.x. In that case, continue with the next step and install python3. You might also need to substitute python3 for python while running commands that use python, such as “python3 --version”, “python3 BoolSim_Windows.py”.
6. To install Python on your Windows machine, follow the steps in this video: https://youtu.be/4Rx_JRkwAjY You may stop at 4:00 in this video once Python is installed. Close the installer and go to step 5. You should be able to see python’s version (3.8 or higher) when you run “python --version” on cmd (as shown in screenshot 1).
7. Next, we install a few required Python packages: numpy, matplotlib, pandas, and jupyter. Run these commands in the command prompt after Python is installed:

a. pip install numpy==1.19.3
b. pip install matplotlib
c. pip install pandas
d. pip install jupyter
8. After the above packages are installed, you may check their version by trying the commands given in screenshot 2 below.
9. Good to go!

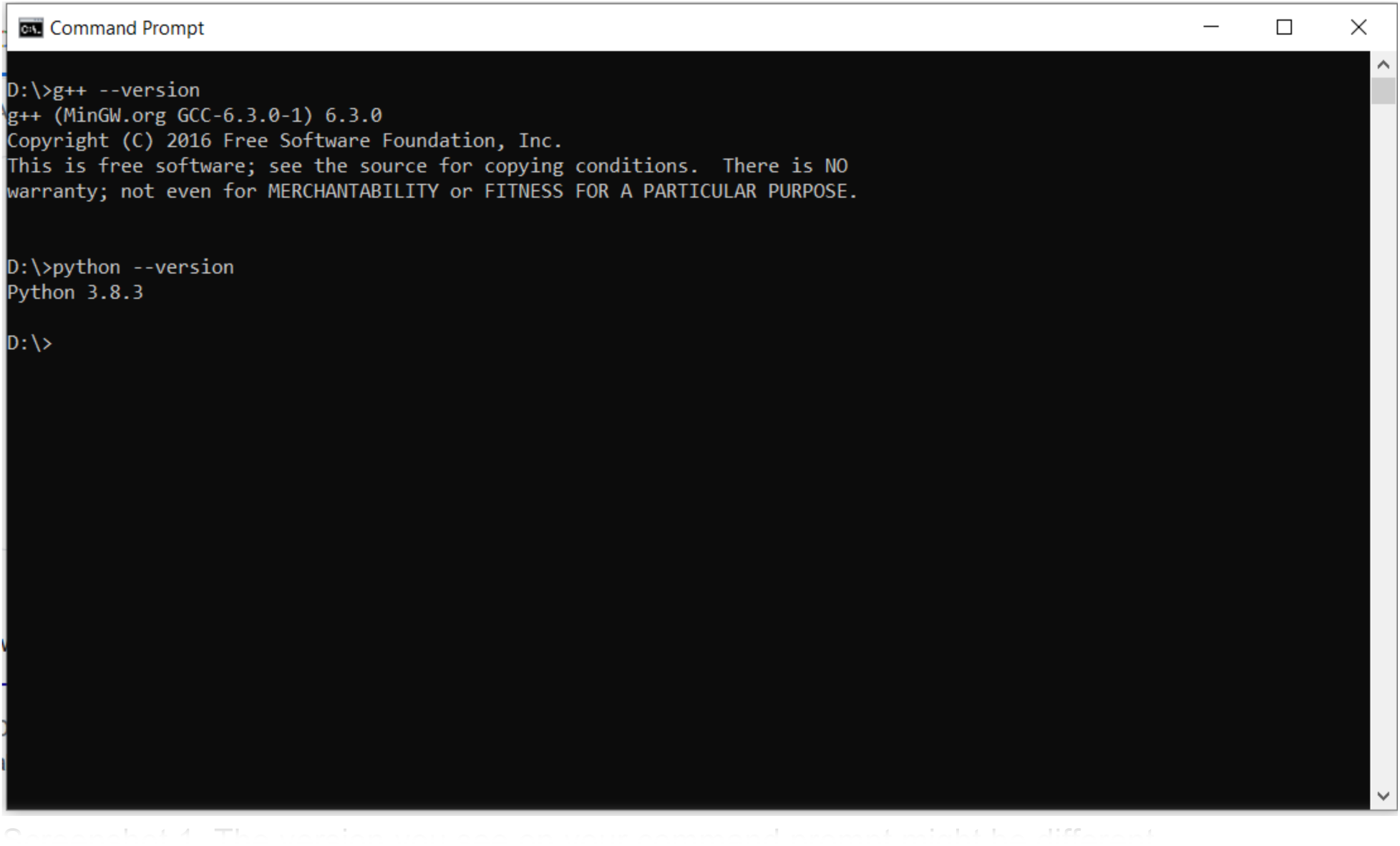
Screenshot 1. The version you see on your command prompt might be different.

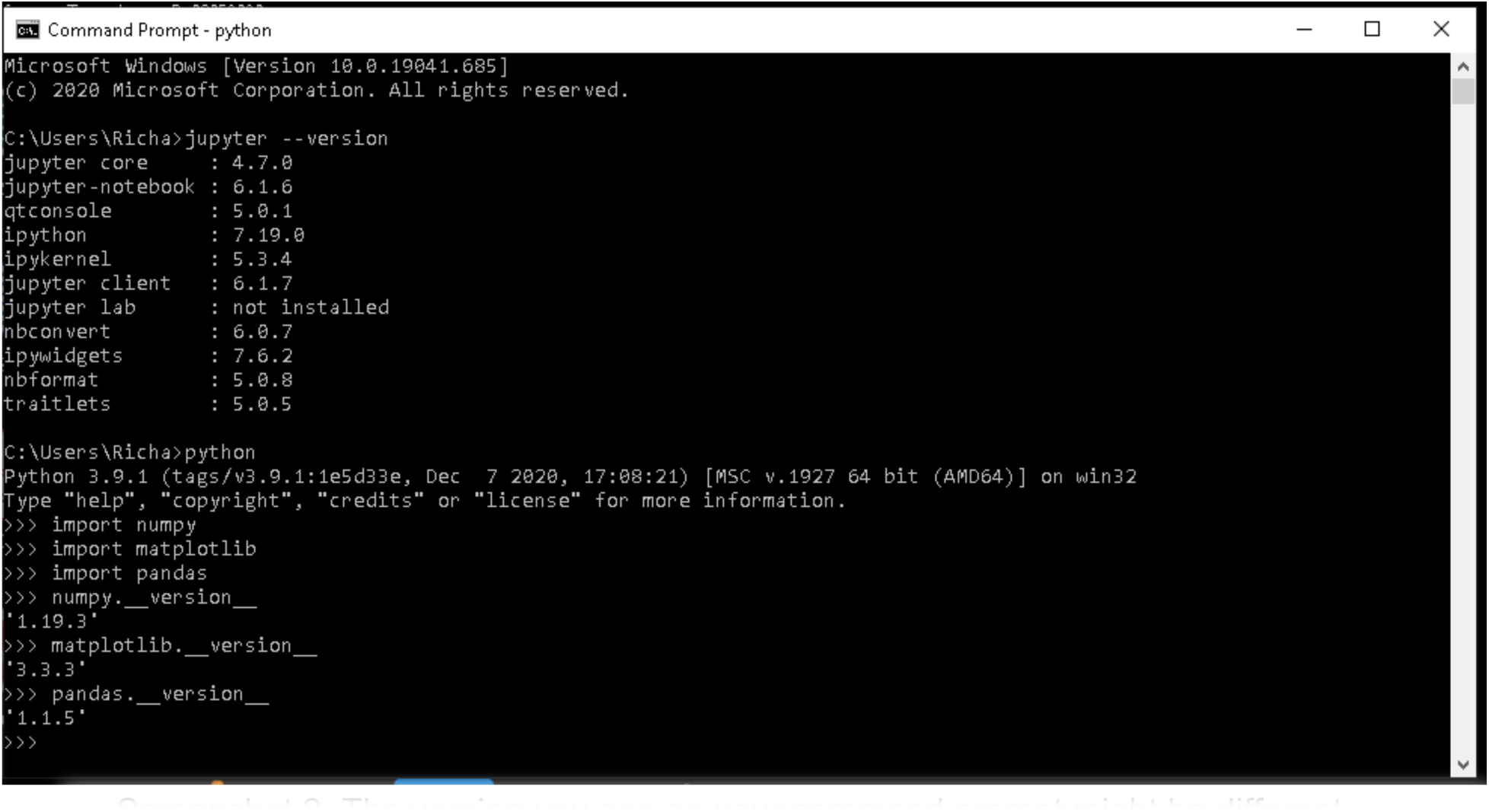
Screenshot 2. The version you see on your command prompt might be different.

### 2. Downloading BoolSim

1. Create a new folder on your computer where you want to run BoolSim. For example, D:\BoolSim_simulation\
2. Visit https://github.com/dyhe-2000/BoolSim-GUI.git or https://github.com/Rappel-lab/BoolSim-GUI to download the package. Click on the green button named Code and then click on Download ZIP in the dropdown. See Screenshot 3.

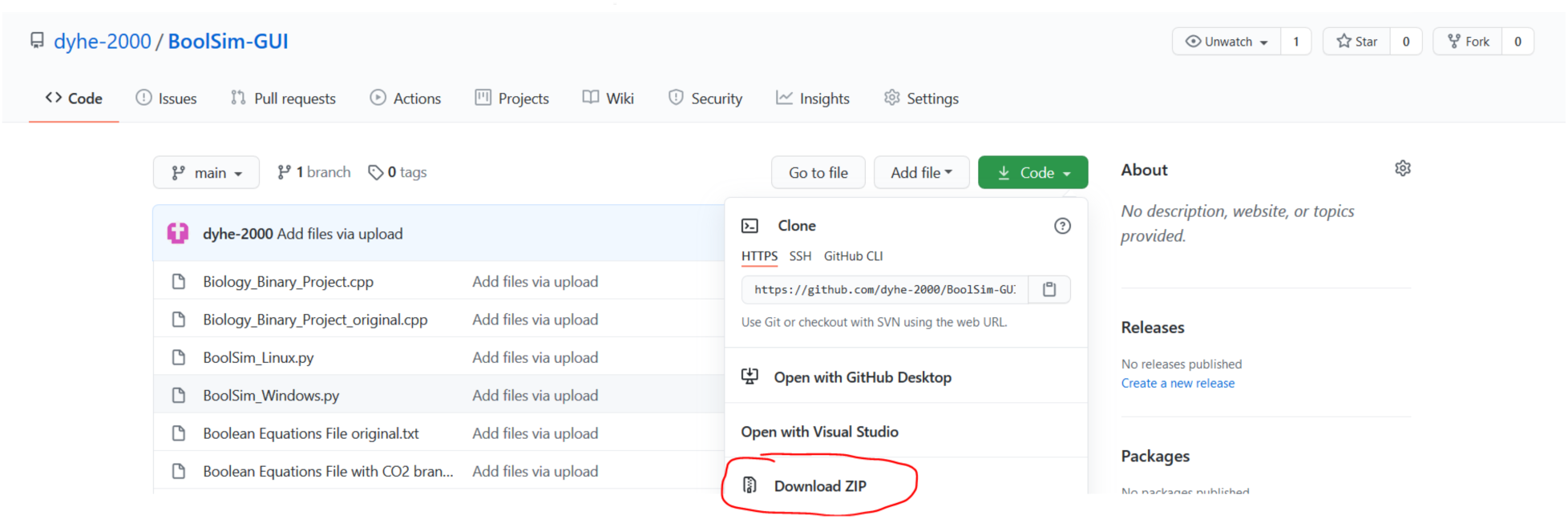
Screenshot 3: Download page of BoolSim from GitHub

3. Save the zip file in the new folder you’ve just created. If the zip file is saved to the Downloads folder by default, cut and paste the zip file into the new folder.
4. Then open the new folder you created in File Explorer. Right-click on the zip file and extract it at the current location. A new folder by the name “BoolSim-GUI-main” is created. Open that folder. It contains all the files needed to run BoolSim.
5. To run BoolSim, we open the command prompt at the current location. To do that, select the address bar in the File Explorer by pressing Alt+D or by right-clicking on the address bar. Once the address is selected (in our example D:\BoolSim_simulation\BoolSim-GUI- main\), replace the address by the command ‘cmd’ in the address bar and hit Enter. A command prompt pops open.
6. In the command prompt, type ‘python BoolSim_Windows.py’ (“python3 BoolSim_Windows.py” to avoid a conflict if python2 is already installed) and hit Enter. A new window will pop up.
7. Click on ‘Agree’ to enter the start page of BoolSim. This is the **Home Page** of BoolSim. You are now ready to use BoolSim! (See Screenshot 4 and the locations of Menu Bar and Home Page options. We will be referring to these often in the following steps).

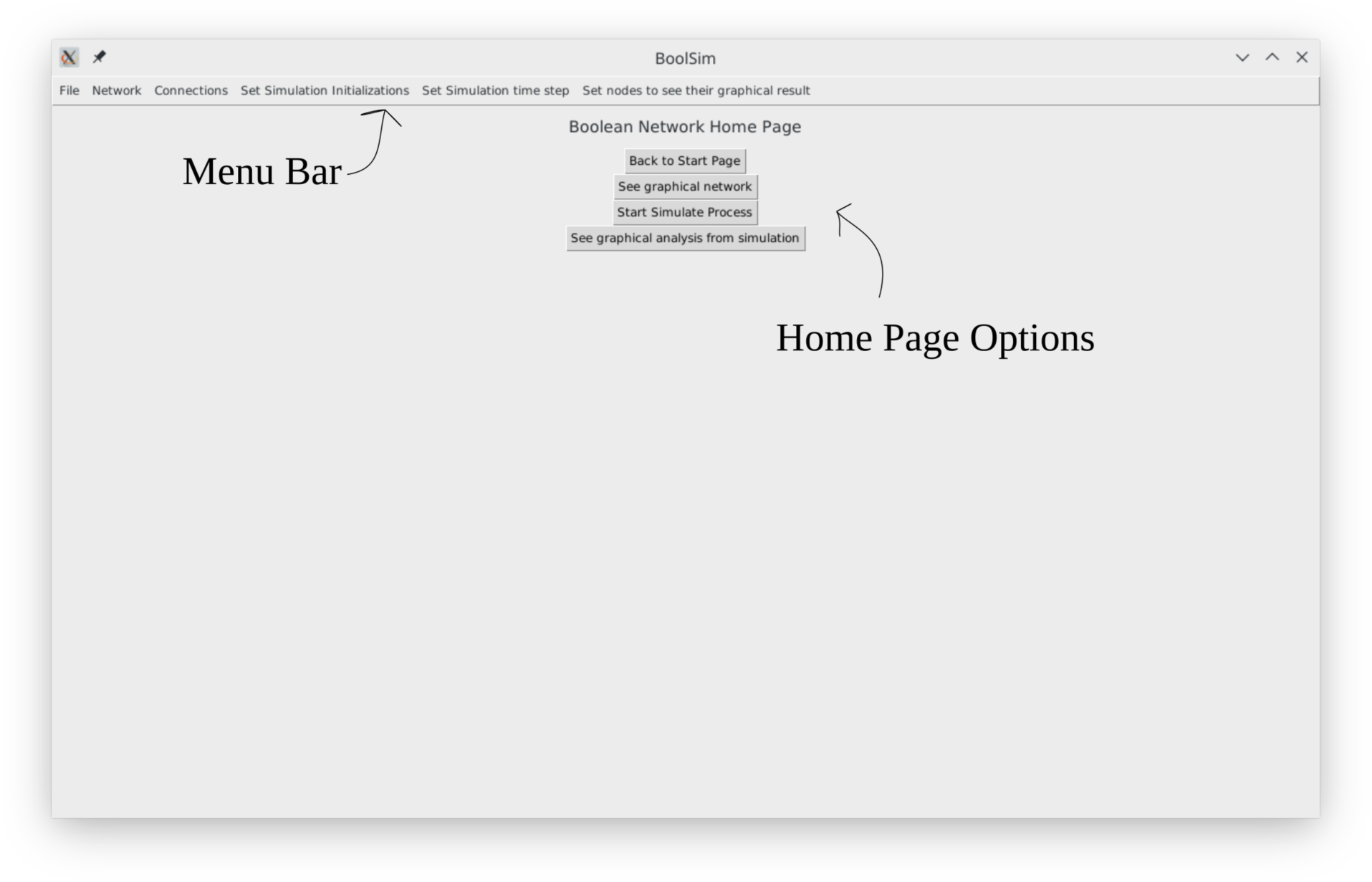
Screenshot 4. Home page of BoolSim

### 3. Running BoolSim

To simulate a Boolean network using BoolSim, you need information on the network. In BoolSim, that information is organized into two files - the first containing the names and initial states of the nodes (if the latter are pre-determined) which we call **node-name-file**, and the second containing the Boolean update equations for each of the nodes defined in node-name- file, which we call **equation-file**. Sections 4.1 and 4.2 describe how to create these files from empirical data. The BoolSim repository contains the files corresponding to three example networks: a “simple network”, the ABA-induced stomatal closure network, and the ABA-CO_2_- induced stomatal closure network (both versions Stomata 2.0 and Stomata 2.1). Refer to Section 4 to locate these files in the current folder.

It is recommended that you try out the simple network first before simulating more complicated networks. The following steps assume that you have located the node-name-file and equation- file for the network you wish to simulate.

To run BoolSim using the files, please proceed with the following steps.

1. To load the nodes of the network into BoolSim,

a. Click Network ➝ Add Node (in the menu bar).
b. Copy-paste the contents of the node-name-file into the text box in the pop-up menu titled ‘Adding Nodes’, and click on the ‘Add’ button.
c. Close the pop-up menu. See screenshot 5.

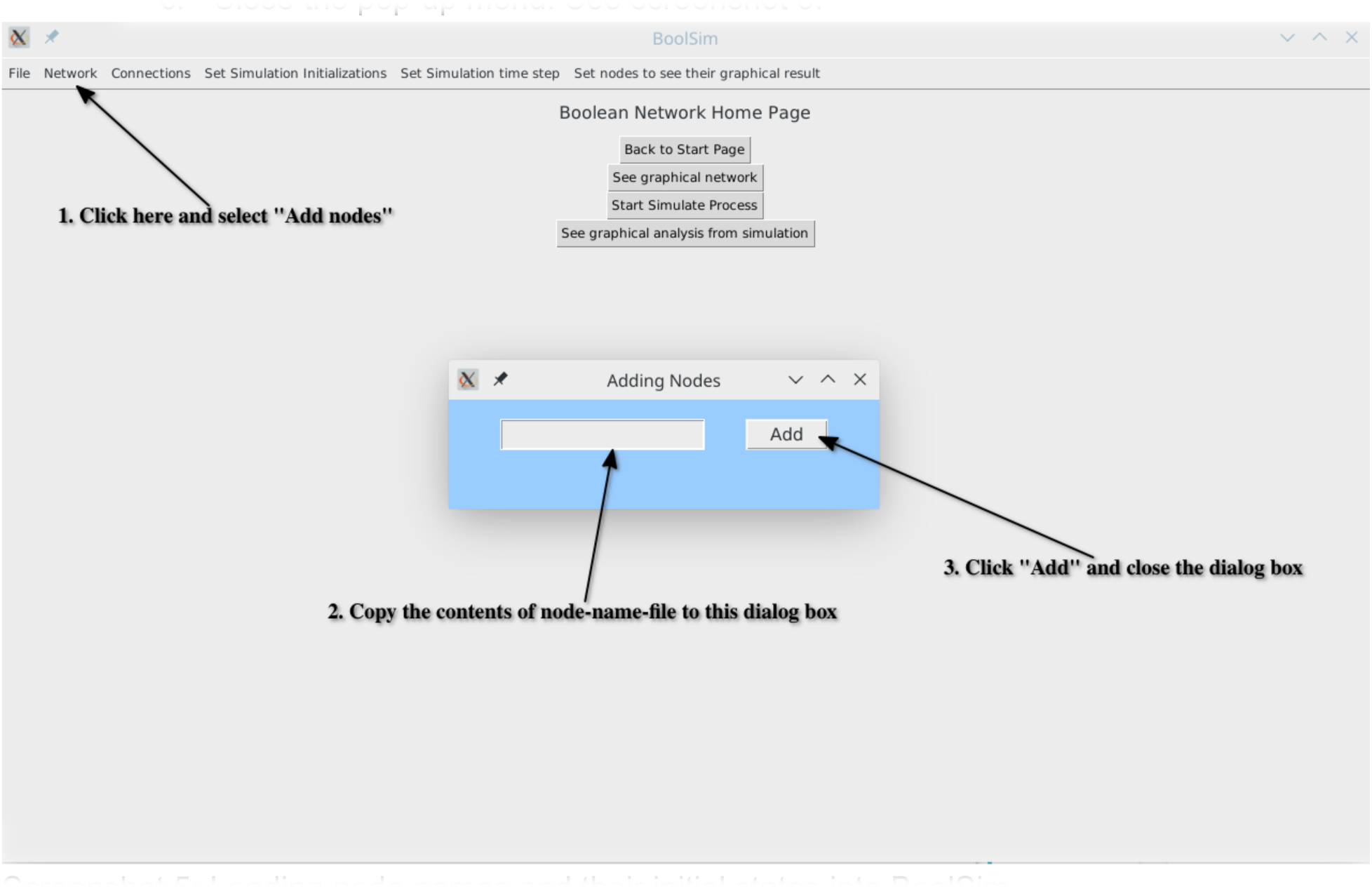
Screenshot 5: Loading node names and their initial states into BoolSim.

A specific example of a node-name-file can be found in the folder Booksim\BoolSim-GUI-main\ sample_data_files \ simple_network_data_files. The node-name-file is called simple_net_node_names.txt and represents the simple network (see also Section 4).

Next, we load the Boolean update equations into BoolSim. BoolSim admits equations written in two formats, word format and index format. The former is the natural way of writing the equations, i.e., with the names of the nodes and as shown in the main text. The latter is the internal representation of the network in BoolSim. Since it is easy to get confused while working with the index format, we do not recommend using it. Nevertheless, we provided the data files in that format.

2. **To load the Boolean equations of the network into BoolSim**, you may use the equations in the word format or the index format. We do not recommend using the Index format.

a. To add equations in the word format, click on Connections ➝ Add Eqns in Word Form. Copy-paste the contents of the equations-file (in word form) into the text box in the pop-up menu titled ‘Adding Eqns’, and click on the ‘Add’ button. Then close the pop-up menu. See screenshot 6.
b. To add equations in the index format, click on Connections ➝Add Eqns in Index Form. Copy-paste the contents of the equations-file (in index form) into the text box in the pop-up menu titled ‘Adding Eqns’, and click on the ‘Add’ button. Then close the pop-up menu.

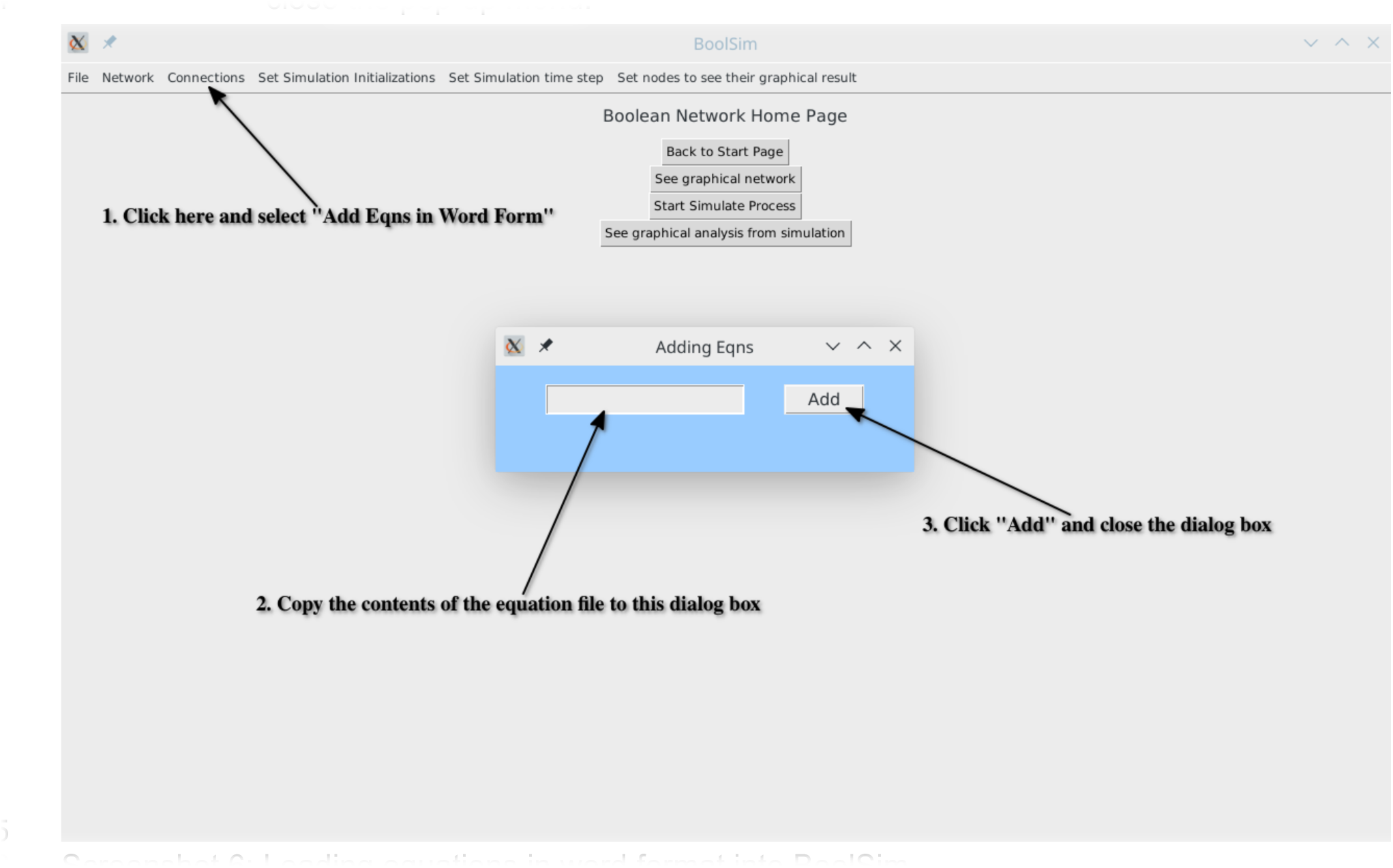
Screenshot 6: Loading equations in word format into BoolSim.

A specific example of a node-name-file can be found in the folder Booksim\BoolSim-GUI-main\ sample_data_files \ simple_network_data_files. The node-name-file is called simple_net_word_eqns.txt and represents the simple network (see also Section 4).

3. To visualize the network, click on the **See Graphical Network** button on the homepage. See screenshot 7. The circles representing the nodes can be moved around by dragging for ease of visualization. Green arrows denote activation, while red arrows denote inhibition. Self-regulation, which can be either positive or negative, is shown by an arrow that is entirely within the node. An arrow begins from the top of the source node to the bottom of the target node. The arrowhead of each arrow is at the target node. Click “Refresh” to reset to the initial view, and click “Back to Home” to return to the homepage. See screenshot 8.

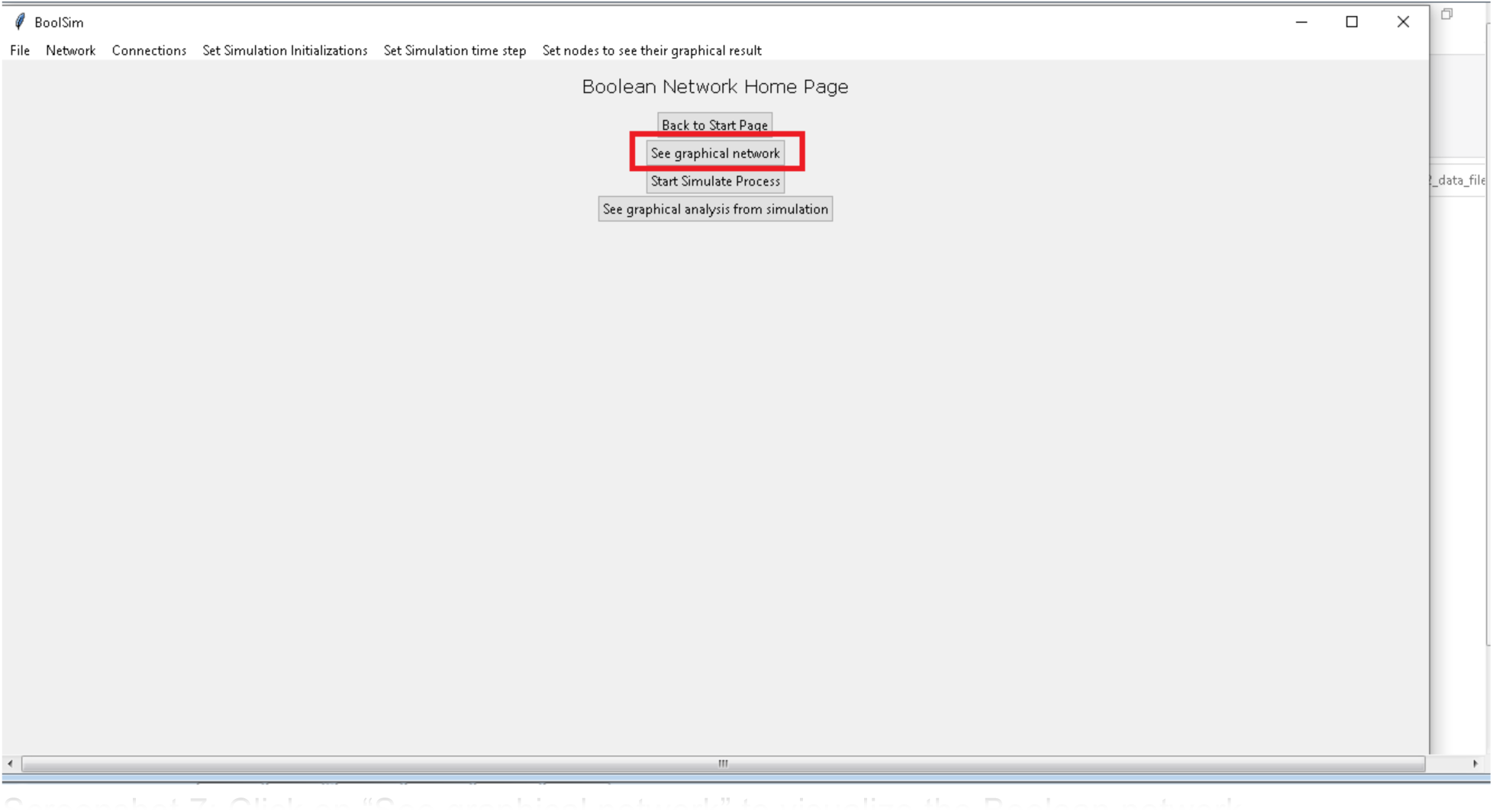
Screenshot 7: Click on “See graphical network” to visualize the Boolean network

4. **To search for upstream and downstream nodes** of the selected node, double-click on the node. The node itself is colored in orange, its upstream nodes are colored in magenta, and the downstream nodes in cyan. See screenshot 8. Right-clicking on or dragging any of the highlighted nodes reverts it to the default yellow.

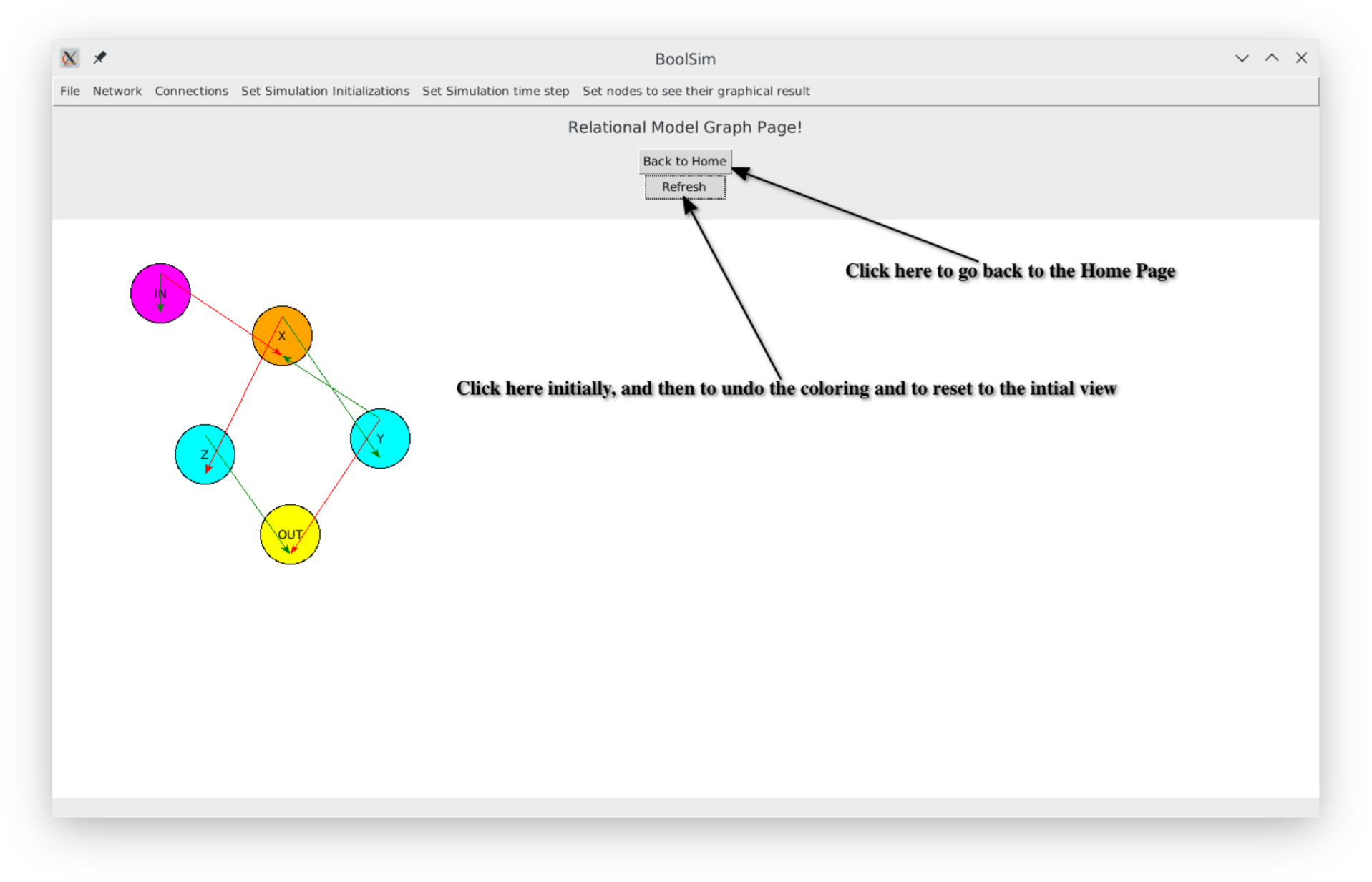
Screenshot 8: Visualizing the “simple” Boolean network

5. **To set the number of initial conditions** before simulating the network, click on the ‘Set Simulation Initializations’ option on the menu bar. You may choose one of the options in the dropdown menu by clicking on them or click on ‘Input arbitrary initializations’ to enter a (different) number into the dialog box. Click on the ‘Add’ button of the dialog box before closing it.
6. **To set the number of time steps,** click on the ‘Set Simulation Time Step’ option on the menu bar. You may choose one of the options in the dropdown menu by clicking on them or click on ‘Input arbitrary time steps’ to enter a (different) number into the dialog box. Click on the ‘Add’ button of the dialog box before closing it.
7. If not at the home page, click “Back to Home”. Otherwise, good to go!
8. **To run the simulation,** click on the ‘Start Simulate Process’ button on the home page. Depending on the complexity of the network and the number of time steps, this may take a few minutes. The button remains inactive while the simulation is running and becomes active once the simulation has been completed. Monitor the progress of the simulation on the command prompt window. At the beginning of each initialization, a message is displayed on the command prompt. See screenshot 9.

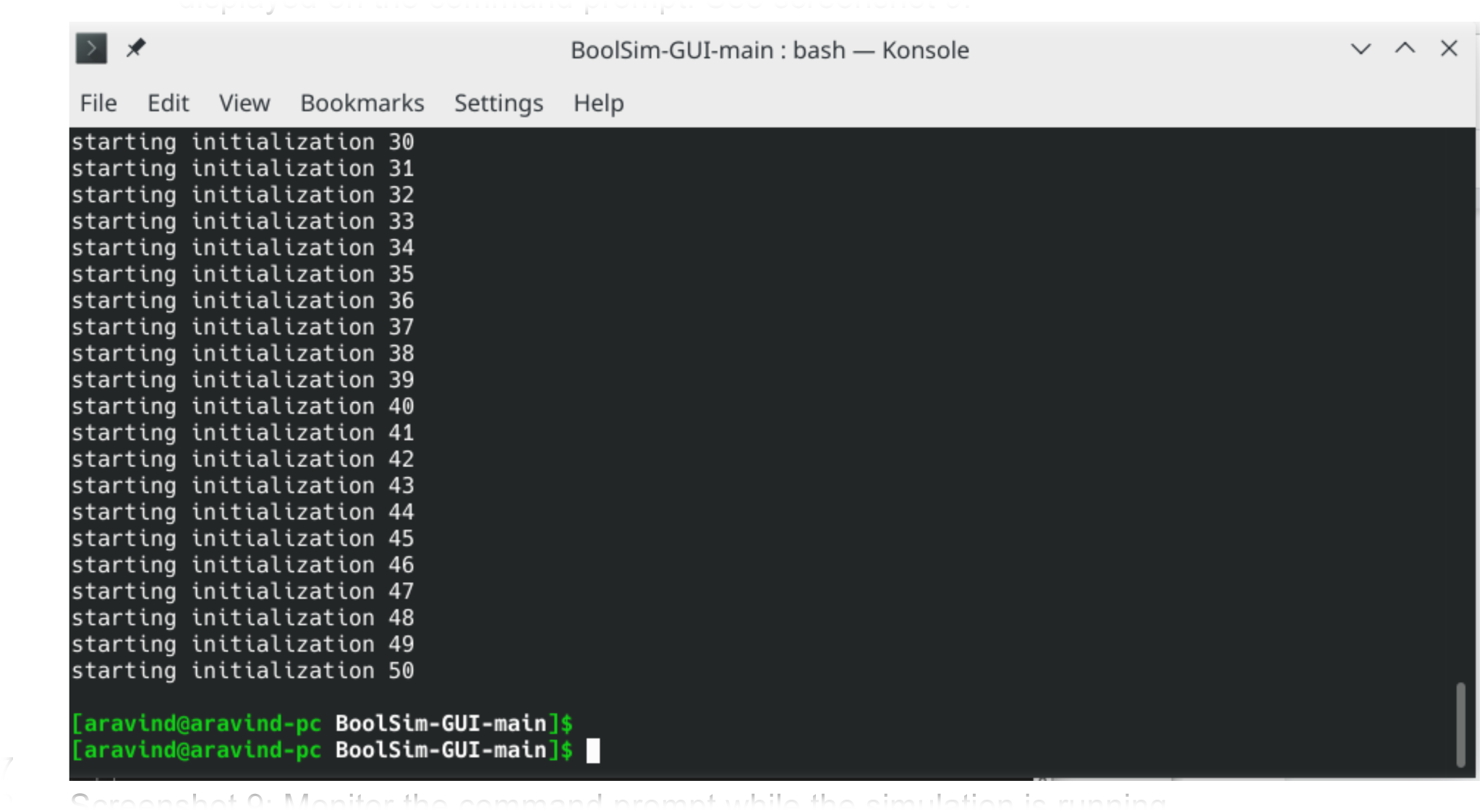
Screenshot 9: Monitor the command prompt while the simulation is running.

After the completion of the simulation, the results are stored in a text file result.txt in the same program folder. Caution: for complex networks, this file can be very large.

9. **To visualize the average activity of a particular node**, click ‘Set nodes to see their graphical result’➝ ‘set node’ on the menu bar and enter the name of the node. If you want to visualize more than one node, a separate procedure is required. For this, you need to open the text file ‘scratch paper.txt’ in the main folder and type each node that you wish to visualize on a new line. Copy all lines and paste it into the ‘set node’ box. For example, if you are simulating the simple network, you may type ‘IN’, ‘X’, ‘Z’ (without quotes) in separate lines in the file ‘scratch paper.txt’. Copy-paste these lines to the text box of the pop-up menu. Click on the ‘Add’ button on the pop-up menu and close it. Click on the ‘See graphical analysis from simulation’ button on the homepage. You will now see a plot and a warning dialog box. Close the warning dialog box and hit Refresh when the plot appears. This last step ensures that the plot only displays the nodes entered in the file. Screenshot 10 shows the activity of the OUT node in the simple network.

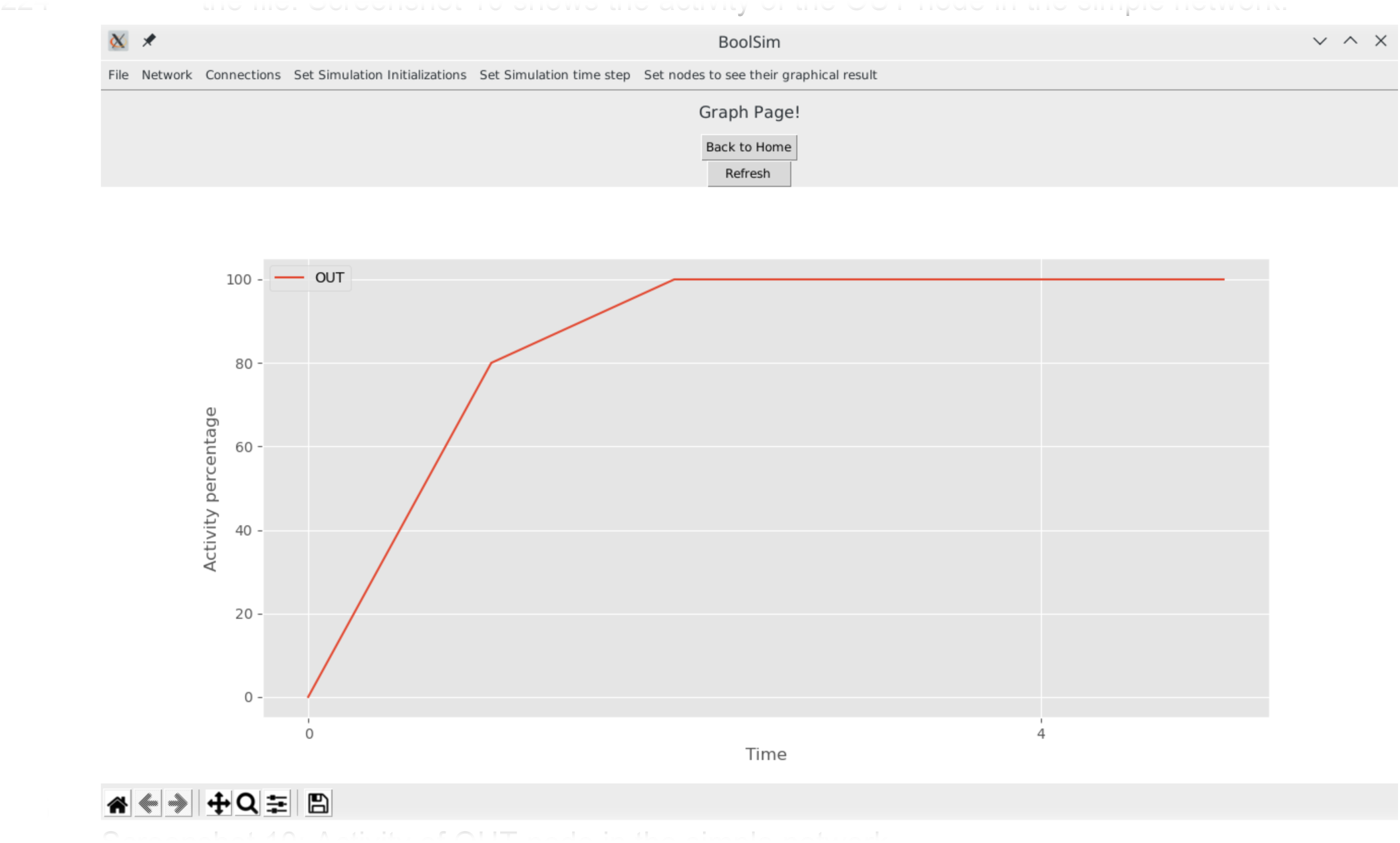
Screenshot 10: Activity of OUT node in the simple network.

10. Click on the ‘Back to Home’ button to get back to the home page of BoolSim in case you want to run a different simulation.

### 4. Preparing the Network

BoolSim comes packaged with text files containing node names and the update equations for a simple network and stomatal closure networks mediated by ABA and ABA+CO_2_. You may first simulate the ‘simple network’ to make sure everything is working fine. To simulate any boolean network on BoolSim, you need a file containing the names and initial states of the nodes (called node-name-file) and a file containing the update equations of the nodes (called equation-file).

The node-name-file and equation-file for the predefined boolean networks are located in the sub-folders as listed below:

1. Folder containing the simple network: sample_data_files \ simple_network_data_files

a. Node-name-file: sample_data_files \ simple_network_data_files \ simple_net_node_names.txt
b. Equation-file: sample_data_files \ simple_network_data_files \ simple_net_word_eqns.txt
2. Folder containing the ABA network: sample_data_files \ ABA_data_files

a. Node-name-file: Node Name and their initial state with ABA.txt
b. Equation-file (you may use one of the two, the word format is recommended):

i. File with equations written in words: Boolean Equations in words with ABA.txt
ii. File with equations written in indices: Boolean Equations in index with ABA.txt
3. Folder containing the ABA-CO_2_ network (Stomata 2.0): sample_data_files \ ABA_CO2_data_files \ stomata2-0

a. Node-name-file: Node Name and their initial state Stomata2-0 .txt
b. Equation-file (you may use one of the two, the word format is recommended):

i. File with equations written in words: Boolean Equations in words Stomata2-0 .txt
ii. File with equations written in indices: Boolean Equations in index Stomata2-0 .txt
4. For Stomata 2.1, refer to sample_data_files \ ABA_CO2_data_files \ stomata2-1. The files are named similarly as in Stomata 2.0

To define your own network, follow the steps below. You may refer to the above data files as templates for the files you create.

#### 4.1 Creating a node-name-file

1. First, create a folder inside sample_data_files to store the files pertaining to your new network. For example, create a folder ‘my_bool_network’ inside sample_data_files.
2. Open Notepad (or your favorite text editor) and save a new file in this directory. For example, save the new file as ‘node-name-file.txt’ in the ‘my_bool_network’ folder.
3. In your node-name-file, type the name of each node of your network in a separate line.
4. If the initial state of a node is to be fixed, instead of being assigned randomly, type the initial state (0 or 1) after a space following the name.
5. In the visualization of the network, if you want a node to be colored in a different color (yellow is the default color), type the color at the end of the line for that node. Available colors are purple, blue, green, yellow, orange, and red. For example, when a node named SLAC1 needs to be colored orange and if its initial state should be 0, you should type SLAC1 0 orange in the line containing SLAC1. When a node named ROS is randomly initialized but needs to be colored blue in the visualization, you should type ROS blue in a new line.
6. The contents of this file will be copied into a dialog box. Save this file before closing.
7. Refer to the node-name-files of the pre-defined networks for examples.

#### 4.2 Creating an equation-file

1. a new text file in Notepad (or your favorite text editor) and save it in the same directory as you did the node-name-file. For example, save this file as ‘equation_file.txt’ in the ‘my_bool_network’ folder.
2. Word format is recommended for the equation-file. The instructions given here are for the word format.
3. Each line in the equation-file contains an update equation for a node. It is recommended to type out the equations in the same order as they appear in node-name-file, though it is not compulsory. The equations should be in the format as described below. There is also a scratch paper.txt for formatting the input to the program. Also, check the existing examples.
4. On the left hand side (LHS), type the name of the node being updated. The names are case-sensitive, so they should be identical to the names in the file for node names.
5. The right hand side (RHS) should be written in the Sum of Products (SoP) form as described in the next section ‘A primer on Boolean Logic’
6. Look at the existing equation-files in word format for examples.

### 5. A primer on Boolean Logic

To specify relationships between variables using Boolean logic, Boolean variables, and functions are used. Boolean variables, like the states of a switch in an electrical circuit, can only take one of two values: OFF (0) and ON (1). In a reaction pathway, the ON state of a variable denotes high activity or concentration while the OFF state denotes low activity or concentration. The relationships between nodes can be described by three basic functions:

- **NOT,** or negation gate: Denoted by NOT or ∼, This operation takes one Boolean variable as input (a unary operator) and returns the *negated*, or complementary state as output, i.e., NOT(1) = 0 and NOT(0) = 1. This operation can also be written compactly as ∼, i.e., NOT(1)=∼1=0.
- **AND,** or conjunction gate: This operation takes two Boolean variables as input (a binary operator) and returns 1 only if both the states are equal to 1. This operation is also called product because the outcome is identical to multiplying the Boolean variables. The compact way of writing this gate is using the symbol &. The truth table, which lists all possible inputs and their respective outputs, for an AND gate is as follows:

| A | B | A AND B |
| --- | --- | --- |
| 0 | 0 | 0 |
| 0 | 1 | 0 |
| 1 | 0 | 0 |
| 1 | 1 | 1 |

- **OR,** or disjunction gate: This operation takes two Boolean variables as input (a binary operator) and returns 1 if at least one of the variables equals 1. This operation is also called sum because the outcome is identical to adding the boolean variables. The short way of writing this gate is using the symbol |. The truth table, which lists all possible inputs and their respective outputs, for the OR gate is as follows:

| A | B | A OR B |
| --- | --- | --- |
| 0 | 0 | 0 |
| 0 | 1 | 1 |
| 1 | 0 | 1 |
| 1 | 1 | 1 |

An arbitrarily complex Boolean function can be composed using the above three operations. A widely used and intuitive way to express boolean equations is the so-called Sum of Products (SoP) form, which is a sum (i.e., series of terms linked by OR logic) of products (i.e., series of terms linked by AND logic and perhaps NOT logic). Consider an example from the ABA-induced stomatal closure Boolean network:

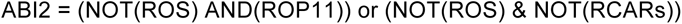

which can be compactly written as

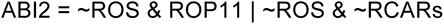

This equation is made of two product terms linked by an OR operator. The simplicity of the SoP form lies in the fact that the equation evaluates to ON (1) if and only if at least one of the product terms evaluates to ON (1). The above equation may also be reported as

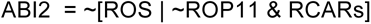

Such a form can be expanded into the SoP form without parentheses using the following identities:

- De Morgan’s laws

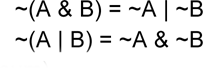

- Distributive law (product over sum)

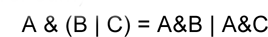

#### 5.1 Deriving Boolean Equations from Empirical Relationships

Boolean equations may be derived from empirical relationships established from experiments, exhaustively measuring the outcomes of all possibilities in a truth table. In the following, we illustrate how to derive boolean equations in the SoP form from a truth table. Consider a node X in a Boolean network, and three nodes A, B, and C upstream of it. Let us say X follows the update rule: X is ON if at least two of its upstream nodes are ON. If we list all possibilities of A, B, and C, and the outcome of X in each case, the truth table looks as follows:

To derive the Boolean update equation for X in SoP form, we gather all the products (connected by ANDs) and add them together (connected by ORs):

**Table 1:** Truth table for X and its three upstream nodes A, B, and C. X is ON if at least three of its upstream nodes are ON.

| A | B | C | X |
|---|---|---|---|
| 0 | 0 | 0 | 0 |
| 0 | 0 | 1 | 0 |
| 0 | 1 | 0 | 0 |
| 0 | 1 | 1 | 1 |
| 1 | 0 | 0 | 0 |
| 1 | 0 | 1 | 1 |
| 1 | 1 | 0 | 1 |
| 1 | 1 | 1 | 1 |

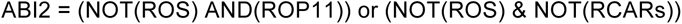

This is the correct Boolean update equation since it considers each case where the outcome is ON separately and exactly once. However, by allowing some redundancy and using basic Boolean logic, the equations in some cases can be simplified and made to look more intuitive. For our example, we can repeat an existing term in the sum without changing the outcome. We may thus write X=(∼A+A)&B&C + A&(∼B+B)&C +A&B&(∼C+C), where we have added the term A&B&C twice. Next, we use the ∼A+A=1, ∼B+B=1, and ∼C+C=1 to obtain X=B&C + A&C +A&B. Finally, replacing each sum with OR, we get

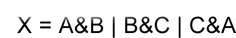

#### 5.2 Analysis of a simple Boolean network

section 2 of the main text described a simple Boolean network with three intermediate nodes and input and output nodes. The network in the main text was compactly written in terms of the symbols &, |, and ∼. In terms of the logical operations AND, OR, and NOT, the equations take on the form

IN = IN

X = Y and (not IN) Y = X

Z = not X

OUT = (not Y) and Z

This illustrative network has the interesting property that while using the asynchronous update scheme and when the input (IN) is absent, the output (OUT) is active 50% of the time. When the input is present, the output is always active. In the former case, the system has an equal chance of reaching one of the two attractors of the system, only one of which leads to an active output node. In the latter case, the system always reaches the (only) attractor which leads to an active output node.

The evolution of the states to the attractor state for this network can be described as follows. In the case without input, (left panel, figure 4 in the main text) there are two attractor states (0,0,1) and (1,1,0); there is no escape from these states. States (0,0,0) and (0,0,1) directly lead to (0,0,1) while (1,1,0) and (1,1,1) directly lead to (1,1,0). Each of the remaining four states on the lattice can reach either attractor with 50% probability. We illustrate this with an example calculation: consider the probability of the state (1,0,1) reaching (0,0,1). There is a direct connection to (0,0,1) which it can take with a probability ⅓. The other path is through (1,0,0) and (0,0,0). (1,0,0) has two exits, along x and y directions. Choosing the x-direction leads to (0,0,0) and hence to (0,0,1). Therefore the total probability to reach (0,0,1) from (1,0,1) is ⅓ + ⅓*½ = ½. Having calculated the probability of reaching each attractor starting from each node, it is easy to see that the probability of reaching (0,0,1) starting from a random node is ½. Note that in an asynchronous update scheme, a node can be updated a second time only after all the nodes are updated once, albeit in any order.

In the case with the input, (right panel, figure 4 in the main text) there is only one attractor (0,0,1), and every state reaches (0,0,1) in a few time steps.

Finally, we contrast the trajectories of states obtained above with those by using synchronous update scheme. In synchronous update scheme, all the nodes are updated together based on the previous state of the system. For example, consider the synchronous update of (1,0,1) in the absence of input (IN = 0) which was examined above for asynchronous update. In one step (1,0,1) evolves to (0,1,0), and then the system oscillates between the two states. Evidently, the system trajectory in synchronous update is not ‘continuous’ in the state space.

For the network at hand, the network has only one attractor (0,0,1) in the presence of input and it reaches it in the utmost two updates. In the absence of output, two of the eight states settle to (0,0,1) in one step, two settles to (1,1,0) in one step, and the remaining four reach the (1,0,1) D (0,1,0) oscillation in the utmost two steps.

### 6. Troubleshooting

Here we list some common errors - and their fixes - that could be encountered while installing various packages and running BoolSim.

1. **Python or g++ are unrecognized**. This happens when g++ and/or Python are not installed on the computer yet. See screenshot 11.

a. You may encounter this error even if you have Python installed in some other way, say with Spyder. It is advisable to do a fresh and complete installation of Python as described in the video referenced in Section 1.

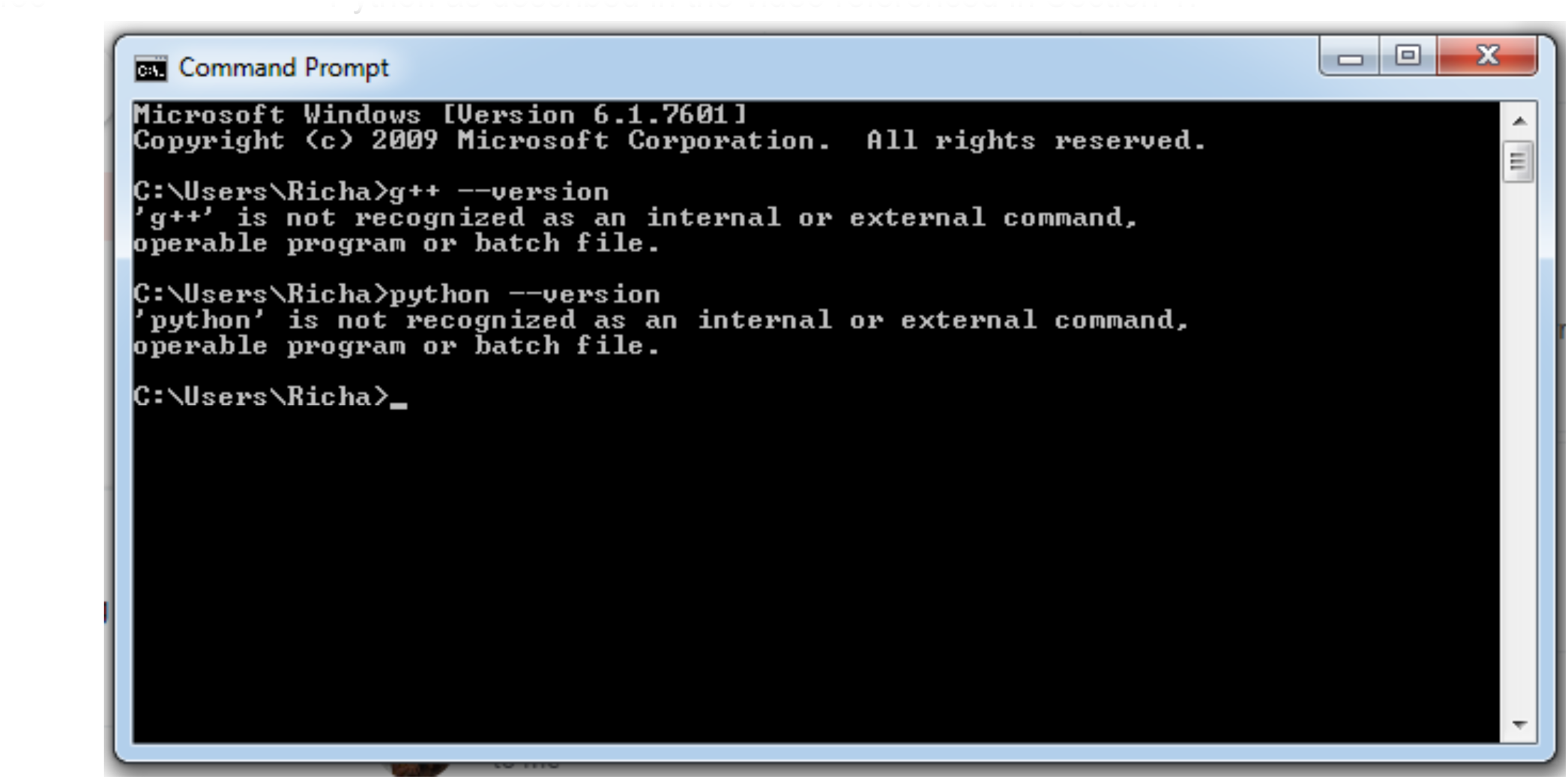
Screenshot 11: g++ and python are not recognized as commands on the command prompt before they are installed on the computer.

2. **Numpy installation fails to pass a sanity check.** This seems to be a common problem with installing Numpy on Windows systems, especially if you tried running the command “pip install numpy”. See screenshot 12. The error would not occur if you run “pip install numpy==1.19.3”, specifying the version number too, as mentioned in the instructions in Section 1.

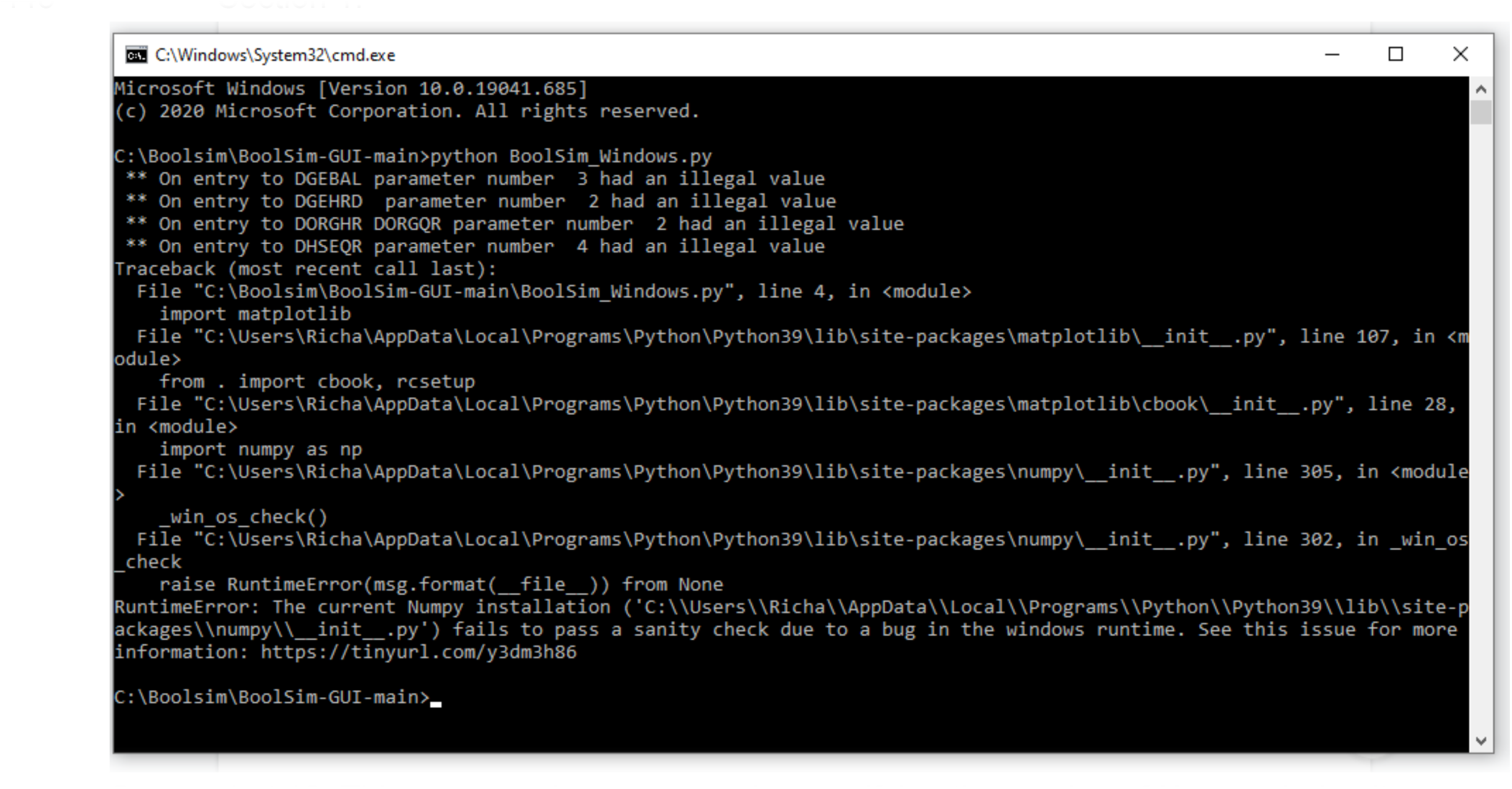
Screenshot 12: This error can be overcome by specifying the version of Numpy during its installation

3. **List out of range.** This error might be encountered when you try to run “python BoolSim_Windows.py”. See Screenshot 13. In this case please check ‘Boolean Equations File’, ‘Word Boolean Equations File’, ‘Node Name and their initial state’ in the folder ‘BoolSim-GUI-main’. These files should be empty. If something is written in these files, then please empty and save them, and try again.

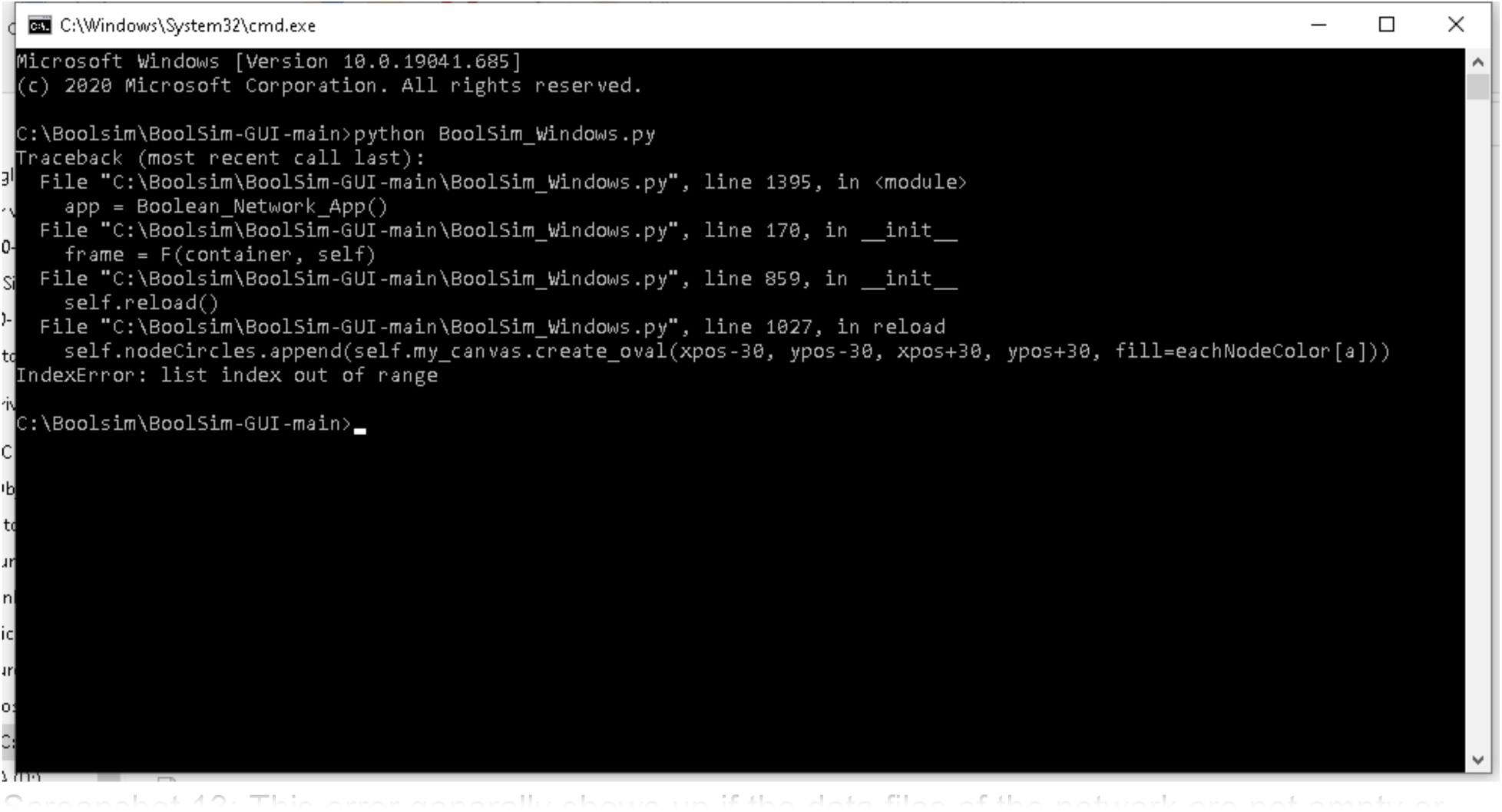
Screenshot 13: This error generally shows up if the data files of the network are not empty or compatible with each other while starting BoolSim.

### 7. Stomata 2.1

As detailed in the main text, our initial attempt Stomata 2.0 to include CO_2_ signaling into the ABA network involved a branch upstream of GHR1. However, the experimental results are inconsistent with the results of Stomata 2.0, which motivated us to seek ways to improve the network. Our experimental results showed that the steady-state conductance level in the absence of ABA is higher for low CO_2_ than for high CO_2_. In the Boolean simulations, we vary the input nodes ABA and CO_2_ between 0 and 1, corresponding to low and high values, respectively. The output is the node Closure, which can be related to stomatal conductance through Conductance=1-Closure. Thus, we want the model to reflect the following experimental observations:

1. Leaves equilibrated at low CO_2_ concentrations have a higher conductance than those equilibrated at high CO_2_ concentrations *prior to* the application of ABA. This should correspond, in the model, to higher conductance for (CO_2_=0, ABA=0) than for (CO_2_=1, ABA=0). Our experimental data suggest that the stomatal conductance for low CO_2_ concentrations is approximately twice as large as for high CO_2_ concentrations.
2. The steady state conductance of the leaves *after* the application of ABA is greater for the lower CO_2_ concentrations than for higher CO_2_ concentrations. This should correspond, in the model, to higher conductance for (CO_2_=0, ABA=1) than for (CO_2_=1, ABA=1). Our experimental data suggest that the stomatal conductance for low CO_2_ is roughly identical to the stomatal conductance for high CO_2_ before the application of ABA.

To translate these experimental findings into a quantitative Boolean state of the output node, we require the model to reproduce the following observations:

ABA=0, CO_2_=0: Closure=0 (Conductance=1)

ABA=0, CO=1: Closure=0.5 (Conductance=0.5)

ABA=1, CO_2_=0: Closure=0.5 (Conductance=0.5)

ABA=1, CO_2_=1: Closure=1 (Conductance=0)

The Closure node in the ABA network model is affected by two nodes, Microtubule and H_2_OEfflux, through the equation Closure = H_2_OEfflux AND Microtubule. To achieve an intermediate level of closure, required for the conditions ABA=0, CO_2_=1 and ABA=1, CO_2_=0, we need one of the two nodes at 100% activity and the other at 50% activity (fluctuating between 0 and 1). For full closure, we need both nodes at 100% while for full conductance we need both nodes at 0. In Stomata 2.1, this is achieved through the following modifications:

1. Ca_2_c = ∼Ca_2_ATPase & (CIS | CaIM) | (ABA&CO_2_)
2. CaIM = ∼ABH1 & (NtSyp121 | MRP5) | ∼ERA1 | Actin | CO_2_
3. Microtubule = TCTP | Microtubule & ABA
4. H_2_OEfflux = AnionEM & PIP21 & KEfflux & ∼Malate | CO_2_

where the modifications are shown in blue. As a reminder, the symbol & represents AND logic and the symbol | represents OR logic.

#### Motivation for these modifications

1. In the original Albert version of the model, Ca_2_c (cytosolic calcium) has 50% activity due to oscillations between Ca_2_c and Ca_2_ATPase, if and only if ABA=1. Since the original network has no CO_2_ input, this is independent of the state of CO_2_. We added input from CO_2_ so that Ca_2_c has 100% activity if both ABA=1 and CO_2_=1. Adding CO_2_ to calcium was motivated by evidence that cytosolic calcium is involved in CO_2_-induced closure.
2. CO_2_ is added to CaIM through an OR gate to ensure that Microtubule=0.5 when CO_2_=1, even in the absence of ABA.
3. In the original ABA network model, Microtubule was always either 0 (if ABA=0) or 1 (if ABA=1), even though Ca_2_c=0.5. This is because of the feedback loop from Microtubule onto itself. To achieve Microtubule=0.5, we make the feedback loop dependent on ABA. As a result, Microtubule=0.5 if ABA=0 and CO_2_=1 but Microtubule=1 when both ABA and CO_2_ are 1.
4. In the original ABA network model, H_2_OEfflux is 0 if ABA is absent, independent of the state of CO_2_. With this modification, H_2_OEfflux=1 when ABA=0 and CO_2_=1.

As a result of these modifications, the network is able to reproduce the experimental results (See Fig. 5 in the main text). It is straightforward to simulate and visualize the network using BoolSim, which can be used to analyze the response:

When ABA=1 and CO_2_=1, H_2_OEfflux = 1 as all the terms are 1. Furthermore, Ca_2_c oscillations are superseded by a sustained activation because of the ABA&CO_2_ term in modification #1. Hence Microtubule is maintained at 1 as well and Closure=1.

When ABA=1 and CO_2_=0, H_2_OEfflux is at 50% since AnionEM is at 50%, and PIP21, KEfflux, and ∼Malate are at 100% activation. Ca_2_c is at 50% (due to oscillations) in the absence of CO_2_ but the positive feedback of Microtubule is activated in the presence of ABA, making it 100% active. Thus, Microtubule=1, H_2_OEfflux=0.5, and Closure=0.5.

When ABA=0 and CO_2_=1, H_2_OEfflux=1 due to the CO_2_ branch implemented in Stomata 2.0 and also present in Stomata 2.1. Similarly, the addition of the CO_2_ branch to CaIM causes Ca_2_c oscillations even in the absence of ABA. Thus, Ca_2_c is at 50% activity and Microtubule=0.5, as the positive feedback is shut down in the absence of ABA. Thus, Microtubule=0.5, H_2_OEfflux=1, and Closure=0.5.

When ABA=0 and CO_2_=0, both H_2_OEfflux and Microtubule are shut down, and there will be no activity of closure. In other words, Microtubule=0, H_2_OEfflux=0, and Closure=0.

## Notes

### Competing Interest Statement

The authors have declared no competing interest.

